# Stress adaptation of free-living microbes generates novel benefits to plant hosts

**DOI:** 10.64898/2026.06.01.729403

**Authors:** Kevin D. Ricks, Kasturi Bhatt, Megan E. Frederickson

## Abstract

Most microbes that live in or on hosts are not obligate symbionts. Instead, they can cycle between host-associated and free-living phases and experience selection in both environments. Consequently, while the benefits that microbes confer to hosts are often assumed to be a product of host selection, they may also evolve independent of the host-microbe interaction, while the microbe is free-living. We investigated this hypothesis by experimentally evolving a beneficial *Allorhizobium* bacterium we had previously isolated from duckweed (*Lemna japonica*), evolving the strain in the absence of any host at high and low salinity and at high and low nitrogen. Inoculating evolved strains back onto the plants, we found microbial adaptation to high-salinity benefited host plants in high-salinity environments, but only when microbes evolved under low-nitrogen conditions. Microbial adaptation to high nitrogen reduced plant growth, in some cases below that of the uninoculated control. In low-nitrogen environments, microbial growth rate predicted plant benefit in matching salinity environments, suggesting the traits underlying microbial salinity adaptation may be associated with the microbial benefits. Whole-genome sequencing of the evolved strains revealed significant genomic shifts between selective treatments, including plasmid variation and point mutations associated with osmotic regulation, although many of the affected genes have unclear functions. The emergence of microbial benefits to hosts as a byproduct of microbial adaptation highlights that these benefits do not depend on host selection. Rather, predicting the evolution of these symbioses may require understanding both abiotic and biotic selective agents acting on key microbial traits mediating the host-microbe interaction.

## INTRODUCTION

All plants and animals are colonized by microorganisms, many of which enhance their host’s growth or fitness [1, 2]. These microbial benefits to the host, ranging from nutrient acquisition to pathogen defense [3, 4], are often assumed to be the product of host selection [1, 5]. While providing these benefits to hosts can be costly to microbes, host-provisioned resources in the form of nutrients, habitats, or other rewards [6] may compensate for these costs, thereby selecting for host growth-promoting microbes [7]. This reciprocal exchange of net benefits can align the fitnesses of partners, as the host’s success directly supports the microbe’s own proliferation [8]. This host-centric perspective, however, often ignores that many host-associated microbes are horizontally transmitted and can spend significant portions of their evolutionary history as free-living organisms [9–11]. While a free-living phase should select against costly cooperative traits [12, 13], beneficial microbial symbionts remain pervasive even in the free-living phase. Consequently, here we ask: can microbial partner quality (i.e., the fitness benefit that a microbe confers to its host) evolve independently of the host? And if so, which abiotic variables in the free-living environment select for higher or lower partner quality when the microbes are host-associated?

We hypothesize that increasing microbial partner quality can be selected for as incidental byproducts of microbial stress adaptation in their free-living phase. In mutualism theory, this is known as a byproduct mutualism [14, 15], because greater cooperation between species evolves as a byproduct of selection for traits that directly benefit one of the participants (i.e., regardless of any reward from the partner). Byproduct mutualisms are a form of indirect genetic effects, where the microbial genes selected upon by the abiotic environment indirectly alter the host’s performance [16]. For example, microbial adaptation to a given abiotic environment may create an incidental fitness alignment between the host and microbe in that environment, as locally-adapted microbes can reach sufficient densities to provide other benefits to the host. Consider, for example, a microbe possessing genes associated with both host growth promotion and its own tolerance to abiotic stress. Alleles encoding microbial stress tolerance might directly benefit the microbe and increase host growth promotion, even without any adaptation at the growth promotion loci, by facilitating large, locally adapted microbial populations. Such byproduct benefits [14, 15] are consistent with previous research showing that plants in a given habitat often receive the greatest benefits from microbes derived from that same habitat [17–20]. However, in most of these studies, microbes adapted to abiotic stress in the presence of hosts (but see [17]), so selection imposed by the host cannot be separated from selection from the abiotic environment. Furthermore, prior work has not identified the genomic basis of microbial stress adaptation, and has tested the effects of only one abiotic variable at a time. It therefore remains unclear whether byproduct benefits depend on specific selective environments.

Indeed, byproduct benefits need not arise through adaptation in every environment. While microbial stress adaptation could incidentally improve host growth promotion under stress, microbial adaptation to changes in resource availability may have different consequences for symbiont partner quality. High resource availability may relax selection on microbial traits involved in nutrient acquisition or metabolite synthesis that benefit the host. For example, in the well-studied legume-rhizobium symbiosis, rhizobia adaptation to high-nitrogen (N) soils reduces their quality as legume mutualists [21, 22]. Consequently selection from both abiotic stressors and resource availability in the free-living environment may shape the evolution of microbial partner quality, potentially in different directions.

Here, we investigated how microbial adaptation during a free-living phase shapes symbiotic quality when host-associated by experimentally evolving microbes under both low and high salinity stress and low and high resource (i.e., N) availability. We studied a previously uncharacterized *Allorhizobium* strain that we isolated on nitrogen-free media directly from wild duckweeds (*Lemna japonica* L.); this strain belongs to a clade that commonly associates with *L. japonica,* but with unknown functional significance [8, 23]. Pilot work showed that this *Allorhizobium* strain significantly increases *L. japonica* growth under both low and high nitrogen conditions, but provides a significantly larger benefit to hosts growing at low nitrogen (Figure S1). *Lemna japonica*-microbe interactions are a tractable system for this work, as individual plants are only a few millimeters in size, and reproduce through clonal budding, with generation times as little as 1–2 days. These traits make *L. japonica* highly amenable to high-throughput experimentation that we can leverage to phenotype microbial impacts on host growth (see Methods below).

We experimentally evolved *Allorhizobium* under fully crossed salinity and nitrogen treatments and then inoculated single ancestral or evolved strains on *L. japonica* to measure their partner quality. Salinity and nitrogen represent two fundamentally different selective forces, a physiological stressor versus an added resource, which in combination we use to understand the evolution of microbial partner quality and byproduct benefits for hosts. As salinity can be a significant stressor for both duckweed and microbes in their freshwater ecosystems [24], comparing high saline- vs. low saline-adapted strains allows us to evaluate the emergence of fitness benefits to both microbes and plants. We hypothesized these benefits may emerge, independent of the selection from the host, as local microbial salinity adaptation would ensure microbial persistence, with continued access for the plant to their microbial partners. In contrast, we hypothesized that added nitrogen may select against cooperative partners by relaxing selection on beneficial microbial traits that are selectively maintained only under nutrient scarcity [25, 26]. By explicitly evolving *Allorhizobium* without a host plant, any shifts in partner quality can be attributed solely to the byproducts of microbial local adaptation to these crossed environments. Moreover, pairing this approach with whole-genome sequencing, we characterized genetic variants that may underlie local adaptation and mutualist quality in *Allorhizobium*. With this approach, we aim to disentangle the environmental drivers of microbial evolution and offer insights that may scale to more complex host-microbiome dynamics.

## MATERIAL & METHODS

### Experimentally evolving bacterial symbionts

#### Selection phase

We used a bacterial strain, *Allorhizobium* strain MF109-1 (a Gram-negative bacterium in the class Alphaproteobacteria, that we had previously isolated from a duckweed population in Toronto, Canada (Lat. 43.68972, Long. -79.41944). The strain was isolated on Burk’s media, a nitrogen-free media [27], though our prior culturing work in this system has frequently isolated closely related strains on several media types. In pilot work (see Supplemental Methods for details), we tested the ancestral *Allorhizobium* strain on axenic *L. japonica* growing in high-and low-N conditions, and found that *Allorhizobium* is especially beneficial to *L. japonica* growing under minimal nitrogen conditions (Figure S1, Table S1). We then evolved this strain in multiple environments in 96-well plates, such that each well had a continuously evolving bacterial population. We modified Burk’s media to create two salinity treatments (addition of 0 g L^-1^ or 5 g L^-1^ of NaCl) and two nitrogen treatments (addition of 0 g L^-1^ and 0.16 g L^-1^ of NH_4_NO_3_), in a fully crossed design. 5 g L^-1^ NaCl is a significant stressor for both the microbes and plants and at the high end, but within the range of, chloride concentrations observed in urban Toronto aquatic habitats due to winter salt runoff (Figure S2), representing a realistic stress for both the plants and microbes. The concentration of the high nitrogen treatment is within the range of many standard high-nutrient bacterial media types, and thus likely provided the necessary nitrogen for robust bacterial growth. We established 10 replicate evolving populations of each treatment, randomly spread across the plate; other wells on the 96-well plate were sterile blanks, or inoculated with bacteria for a separate evolution experiment. Plates were maintained in a shaking incubator at 25°C. We maintained experimentally evolving populations for 90 days, transferring 10 µL every 2 days from each well to a corresponding well in a new plate with fresh media with the same salinity and nitrogen concentrations.

#### Isolation

Following the selection phase, we isolated a single strain from each evolving population by streaking 10 µL from each well onto yeast-mannitol (YM) plates. While all media types selectively filter bacterial genotypes, we used YM for this step instead of the nitrogen-free Burk’s media we originally used to isolate the ancestral strain because Burk’s media would be a strong filter with respect to the capacity to grow without nitrogen. In contrast, YM is a general resource-rich environment we presumed allowed many bacterial phenotypes to persist. We picked a single colony from each population, and serially plated to purify. Isolating a single colony per replicate ignores within-population strain diversity, but maximizes the number of independently evolving lineages we can test on plants and facilitates linking genomic variants to microbial and plant growth rate in the cross-inoculation experiment (described below). Of the 40 initial populations, three yielded no bacterial growth on YM, all from the high-salinity, high-nitrogen treatment. We speculate this must have been due to a failed transfer at some point over the selection phase. All evolved strains had 16S rRNA sequences that matched the 16S rRNA sequence of the *Allorhizobium* ancestor.

### Characterizing impacts of selection on evolved strains’ partner quality

#### Microbial impact on plant growth

We tested how the ancestral and derived *Allorhizobium* strains impacted plant growth through controlled inoculations onto *L. japonica* plants at both low- and high-salinity. Our previous work describes our methods in high-throughput phenotyping to characterize *L. japonica* growth [28]. Briefly, using a duckweed population we had previously collected from a ravine in Toronto (Lat. 43.68972, Long. -79.41944), we bleach-sterilized individual duckweed fronds (see [29] for details). We moved individual sterile fronds into sterilized Appenroth media and sealed jars, placing these in growth chambers. Appenroth media mimics the nutrient environment of pondwater, duckweeds’ natural habitat [30], and is an ideal media for duckweed growth. Following 2 weeks of growth, populations in each jar reached ∼2000 fronds. We validated sterility through streaking media and plant tissue on general-purpose agar media, including LB and YMA. From these source populations, we then aseptically moved single fronds using sterilized cotton swabs into individual wells in sterile 24-well plates filled with 2 mL of sterile, nitrogen-free Appenroth media with 0 or 5 gL^-1^ NaCl added (i.e., the same NaCl concentration as in the selection phase of the evolution experiment). These experiments were conducted specifically under low nitrogen because the ancestral *Allorhizobium* strain is most beneficial to *L. japonica* in low nitrogen (Figure S1). We established 5 technical replicate wells in each salinity treatment for each of the 37 evolved strains. Each evolved strain was a clonal isolate from a single colony, and represented a biological replicate for their selection treatment. Evolved strains were grown in YM media for two days at 30°C on a shaking incubator, pelleted, and resuspended in nitrogen-free Appenroth media. We pipetted suspended strains into wells to bring the optical density (OD_600_) of the solution to 0.015 (∼1.2 x 10^7^ cells). We additionally included 20 technical replicates of two control treatments in both salinity treatments: wells with media only (uninoculated, sterile), and wells inoculated with the ancestral strain. As the evolved treatments include numerous biological and technical replicates, to achieve comparable statistical power in these controls we increased the technical replication of the control relative to the other treatments. Across all treatment combinations, this resulted in a total of 450 experimental units (37 strains x 2 salinity levels x 5 technical replicates + 2 controls x 2 salinity levels x 20 technical replicates).

We sealed each plate with a Breathe-Easy sealing membrane (Sigma-Aldrich) and placed plates in a growth chamber where we could continuously measure plant growth. Light intensity was set to 250 µmol/s/m^2^, with a 16h/8h day/night schedule fluctuating between 22°C and 18°C . Plates were placed on a clear acrylic table within the growth chamber and a custom imaging system with four cameras attached to a Raspberry Pi board captured images of each plate 3 times daily. We grew plants for 14 days. With short duckweed generations, this experiment length allowed for numerous plant reproductive events. We processed each image using custom computer vision software that measured plant area in each well (details on both image capture and processing described elsewhere [28]). As duckweed primarily reproduces through asexual budding, area is a proxy for fitness.

#### Microbial growth rate under variable salinity and nitrogen

We also measured microbial growth, independent of the plant, across the interactive salinity and nitrogen environments. Similar to the above methods, we inoculated the ancestral and derived strains suspended in sterile PBS into 96-well plates filled with either low or high salinity and either low or high nitrogen Burk’s media, compositionally identical to the media we used in the original selection phase. Inoculations brought the initial cell concentration of each well to 0.005 OD_600_ (∼4 x 10^6^ cells). We placed plates in a plate reader (CLARIOstar Plus, BMG Labtech) for 48 h, set to a temperature of 25°C, with 30 s of shaking every 30 min, following which OD_600_ was measured. Due to limited space on a single plate, we conducted this growth assay across multiple temporal blocks, replicating each strain, nitrogen, and salinity combination 3 times.

#### Microbial genomics

We used whole genome sequencing to identify variants that may underlie specific phenotypic traits in our evolved strains (see Supplemental Methods for details on culturing, library preparation, and assembly). Briefly, from each strain we extracted genomic DNA using PureLink Genomic kits (Life Technologies), followed by short-read sequencing (Illumina NextSeq2000, 150 bp paired-end, 75–300x coverage per strain) of all ancestral and evolved strains, and long-read sequencing (Oxford Nanopore R10.4.1, 150x coverage) of the ancestral strain.

To build a high-quality reference for the ancestral *Allorhizobium* strain, we generated a *de novo* closed genome assembly using the Trycycler workflow [30] with our long-read data, and subsequently polished the assembly with Polypolish and Pypolca [31, 32] using our short reads. We characterized functions from this ancestral reference assembly by first predicting protein-coding genes using Prodigal [33], followed by functional annotation using EggNOG to generate putative gene annotations and assign genes to broad functional categories (Clusters of Orthologous Genes, COGs) [34]. However, because many genes could not be annotated using this method, we additionally passed gene sequences through the InterProScan pipeline [35], extracting GO terms for each gene. While GO terms are general functional categories rather than specific functions, this approach can provide some classification for the majority of genes.

To explore functions by which the ancestral strain might affect plant growth, we annotated predicted proteins with PGPg_finder [36], which assigns gene functions through DIAMOND alignment [37] against the curated PLaBAse (PLant-associated BActeria) database. As this pipeline can also return universal, housekeeping bacterial functions, we restricted annotations to those with direct links to plant performance (see Data S1 for full list). These include functions involved in nutrient acquisition (inorganic phosphate solubilization, nitrogen fixation, and siderophore production), plant hormone production (IAA, GABA, and cytokinin), vitamin biosynthesis (riboflavin, thiamin, and pyridoxine), and stress alleviation (EPS production). Given that we isolated the *Allorhizobium* strain on N-free media and it was most beneficial to *L. japonica* at low N (Figure S1), we also searched the ancestral genome for homologs of genes known to be involved in nitrogen fixation. Specifically, we used HMMER [38] with Interpro protein family profiles for *nifH*, *nifD*/*nifK*, *nifE*, *nifN*, *nifB*, *nifA*, *nifV*, *nifS*, *nifU*, and *fixABCX*. We included only hits with E-values < 1 x 10^-30^,

To identify *de novo* variants introduced in the evolved strains, we mapped the short-read sequences from each evolved strain to our ancestral reference assembly using both the breseq and GATK variant calling pipelines [39, 40], each with their own strengths. We used the breseq pipeline, which was specifically designed for bacterial experimental evolution studies, to identify large insertions and deletions and the GATK pipeline to identify single nucleotide polymorphisms (SNPs). We used multiple methods to identify novel (compared to the ancestor) genomic variants in the experimental treatments that may explain the observed microbial traits. We characterized if the distributions of variants were structured by the selection treatment, assessing if genomic distances between evolved strains are explained by treatment. We additionally assessed if specific SNPs were in regions of genetic differentiation between treatments by characterizing differentiation in SNP enrichment in the genome. We then evaluated how plasmid loss was structured across treatments, and the functional capacity of these plasmids. These approaches assume that microbial traits will convergently evolve using the same genomic pathways. However, mutations across disparate genes, but with similar functions, could produce similar phenotypes [41]. To potentially capture this variation, we additionally assessed if the selective treatments could explain SNP distribution based on their functional annotations. Additional details on these analyses are in the Supplemental Methods.

### Data analysis

#### Microbial impact on plant growth

We used linear mixed models to characterize shifts in bacterial impacts on plant growth. We modeled plant area estimates from all 42 images of each plate captured over the 14-day experiment. Within our model, we included as fixed effects: the timepoint at which the image was taken, the contemporary salinity environment for the plant (low or high), the historic salinity environment for the inoculated strain (low or high), the historic nitrogen environment for the inoculated strain (low or high), and all two-way interactions between these terms. We additionally included the initial area of plants within each well. As random effects, we included plate and well ID, as each well was measured repeatedly and therefore non-independent. We also included strain ID as a random effect, nesting the 5 technical replicates within their respective independently evolved lineage. We constructed models and evaluated the fixed effects using the lme4 and lmerTest packages [42, 43] in the R statistical environment. We extracted model coefficients and their standard errors, specifically focusing on interactions involving the timepoint term. This timepoint term models the change in plant area over the course of the experiment, representing a plant growth rate; interactions with this term consequently represent how various treatments influence the plant growth rate. Increasing plant growth rate means higher microbial partner quality. While these models describe how plant growth changes in response to the selection treatments we imposed on the bacteria, we built additional models to estimate individual strain effects on plant growth. These models were similar to above, but replaced the historic nitrogen and salinity treatments with the individual strain identity.

#### Microbial growth rate under variable salinity and nitrogen

We estimated microbial growth in the nitrogen and salinity environments by generating strain-level estimates for maximum microbial growth rates in these environments. For bacteria that reproduce by binary fission, microbial growth and fitness and one and the same. As microbial growth curves may frequently not fit parametric growth models, we estimated growth rates by using a smoothing spline, passing each replicate through the fit_spline function from the growthrates R package [44]. As our main interest was in how strain-level fitness variation was associated with plant benefits, we built mixed models to estimate growth rate for each strain, using strain ID as the fixed effect and plate run as a random effect.

## RESULTS

### Bacterial salinity history enhances, while nitrogen degrades, mutualist partner quality

Experimental evolution of *Allorhizobium* without plant hosts significantly altered the benefits this bacterium provides to plants (Figures 1 and S3, Table S2). In our models, differences in plant growth rates among treatments are represented by interactions between treatment and timepoint terms. While the contemporary high-salinity environment predictably slowed overall plant growth by 21% (significant timepoint × contemporary salinity interaction in Table S2), this effect was contingent on microbial evolutionary history (i.e., there were significant timepoint x contemporary salinity x historical salinity/nitrogen treatments, as well as a significant timepoint x contemporary salinity x historical salinity x historical nitrogen interaction term, see Table S2). In the contemporary high-salinity environment, while there was considerable variation among evolved strains (Figure S3), inoculation with strains evolved in historical high salinity on average increased plant growth rates by 18% compared to strains adapted to low salinity, but only when strains evolved at low nitrogen (Figure 1B). Bacterial strains evolved under high nitrogen and high salinity did not differ in quality from strains adapted to low salinity (Figure 1B). In other words, by adapting to high salinity, *Allorhizobium* became a more beneficial symbiont for plants growing at high salinity, but only when the strains also evolved at low nitrogen.

**Figure 1:**
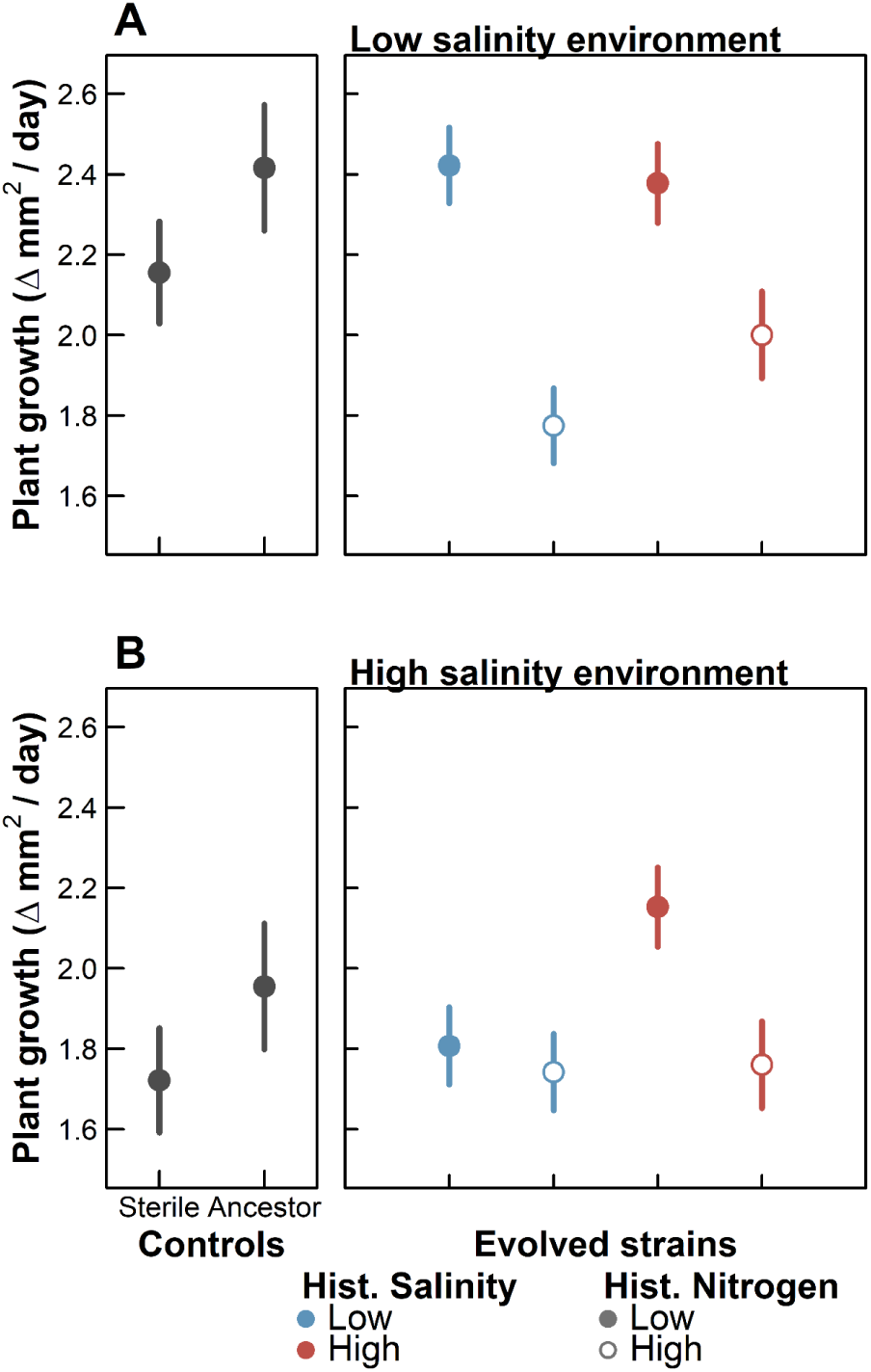
Evolutionary history of the experimentally evolved *Allorhizobium* strains impacts plant growth in both low (**A**) and high salinity (**B**) environments. Dots are estimated marginal means of plant growth (mm^2^/day) when plants were inoculated with a sterile (i.e., microbe-free) control or the ancestral *Allorhizobium* (left panels, black symbols) or with *Allorhizobium* strains that evolved at low (blue symbols) or high (red symbols) salinity and low (closed symbols) or high (open symbols) nitrogen (right panels). Error bars are 95% confidence intervals generated from standard errors. Full model results are in Table S3 and individual strain estimates are in **Figure S3.**

In contrast, bacterial adaptation to high nitrogen consistently degraded the microbe’s quality as a mutualist. When tested under low-salinity conditions, plants inoculated with high-nitrogen adapted bacterial strains grew 34% slower than those inoculated with low-nitrogen adapted strains (timepoint × historic nitrogen interaction; Table S2, Figure 1A). Moreover, plants inoculated with these high-nitrogen adapted strains grew 12% slower than sterile, uninoculated controls, suggesting strains may have evolved mechanisms that harm their hosts. Thus, bacterial adaptation to high salinity and high nitrogen had opposite effects on their quality as plant symbionts (Figure 1); bacterial adaptation to high salinity benefitted plants under high salinity, but only if bacteria evolved under low nitrogen (Figure 1B), while bacterial adaptation to high nitrogen reduced their symbiotic quality, especially when tested at low salinity (Figure 1A).

In evaluating the individual evolved strain impacts on plant growth, there was no evidence for tradeoffs among strains in the microbial partner quality across the two salinity environments (Figure S4). In fact, there was a slight, though non-significant, positive correlation in partner quality between environments. This correlation, however, appeared to be largely driven by strains adapted to high-nitrogen environments having low quality in both salinity environments.

### Microbes adapted to their local salinity and nitrogen environments

Experimental evolution of *Allorhizobium* strains significantly altered bacterial growth rates across the environments, and was largely consistent with local adaptation (Figure 2, S5, Table S3). Strains generally had the highest mean growth rate in the contemporary environment that matched the historical environment in which they evolved. There was also evidence for some tradeoffs in microbial adaptation between environments (Figure S6). Strains with the highest growth rates in high nitrogen conditions generally had among the lowest growth rates in low nitrogen, although only at low salinity.

**Figure 2:**
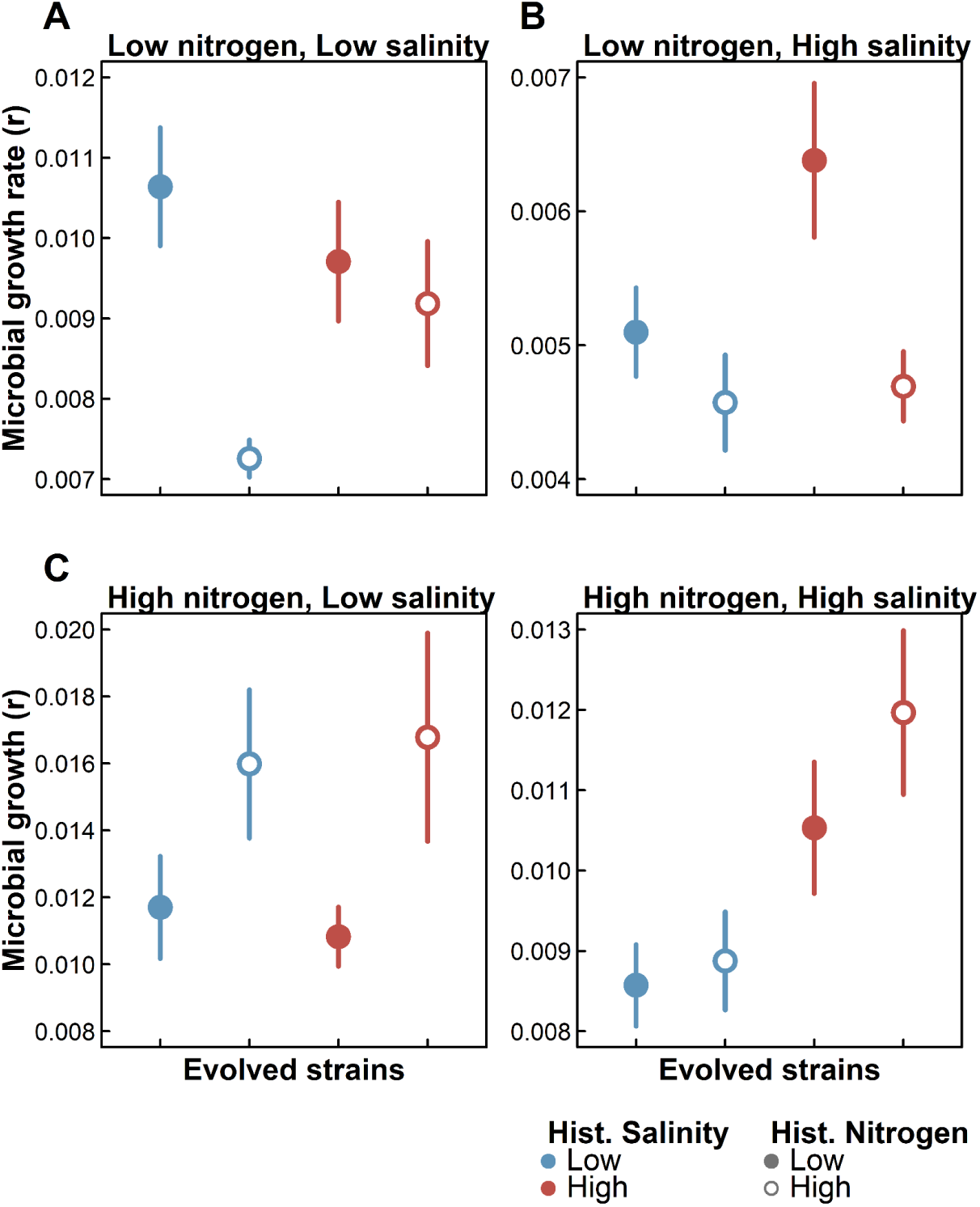
Evolutionary history of the experimentally evolved *Allorhizobium* populations impacts their growth rate across environments. Dots are estimated marginal means of microbial growth (r, calculated from growth curves for each strain) under (**A**) low nitrogen and low salinity, (**B**) low nitrogen and high salinity, (**C**) high nitrogen and low salinity, and (**D**) high nitrogen and high salinity for *Allorhizobium* strains that evolved at low (blue symbols) or high (red symbols) salinity and low (closed symbols) or high (open symbols) nitrogen. Error bars are 95% confidence intervals generated from standard errors. Note the different y-axis scales among panels. Full model results are in Table S4 and individual strain estimates are in Figure S5.

Finally, faster-growing microbes had higher partner quality, but only at low nitrogen. Strain growth rate in low-nitrogen conditions was highly correlated with the strain’s impact on plant growth (Figure 3) when matching the salinity environment. For example, strain growth rate under low nitrogen and high salinity was positively correlated with its impact on plant growth in that same environment. Strain growth rates in non-matching environments were largely poor predictors of plant growth (Figure S7).

**Figure 3:**
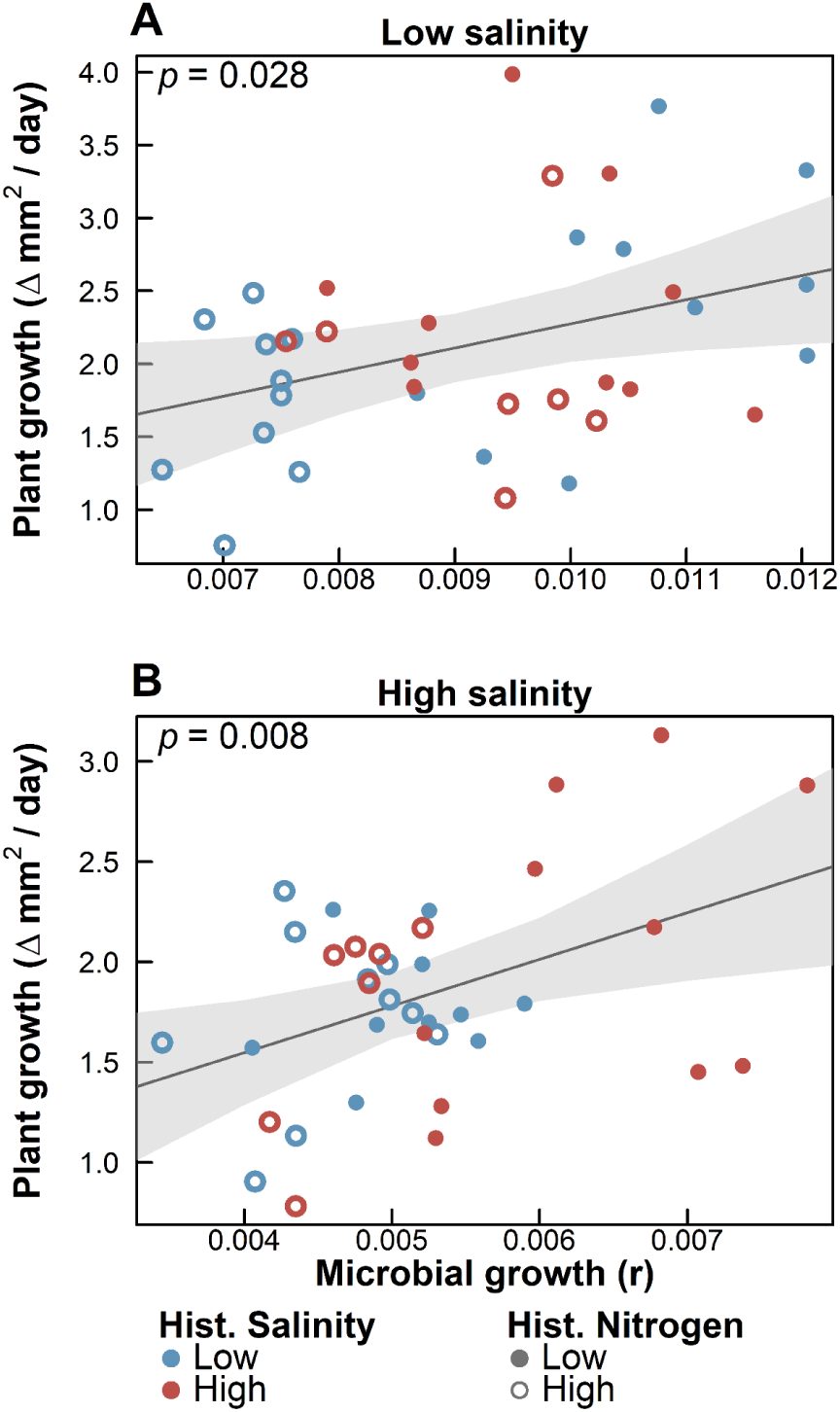
Microbial benefits provisioned to the host correlated with their growth in both low (**A**) and high (**B**) saline environments. Dots are mean plant growth rate (y-axis) and mean microbial growth rate (x-axis) for *Allorhizobium* strains that evolved at low (blue symbols) or high (red symbols) salinity and low (closed symbols) or high (open symbols) nitrogen. Plant growth and microbial growth are the same as those presented in Figures 1 and 2 respectively, though broken down by individual strain. Correlations among bacterial growth across all environments are in Figure S5. The solid line represents the predicted output from a linear regression correlating these traits, and the estimated *p* value in the top left corner of each panel. Note the separate y-axis scales in top and bottom panels.

### Genomic characterization of strains

Our assembly produced five major, fully circular, unfragmented contigs (details in Table S4). Based on size, we assume the largest replicon is the main bacterial chromosome and the other four are plasmids (i.e., small, circular extrachromosomal DNA elements that can be highly mobile, termed plasmids A–D). This assembly was highly similar to an *Allorhizobium* strain found on an alga in the UK (BioSample: SAMEA114008338).

Comparing predicted proteins against a curated database of plant-growth promoting genes identified multiple candidate plant growth promotion genes, including siderophore production, phosphate solubilization, plant hormone production, vitamin biosynthesis, and EPS production (Table S5). Most of these genes were found on the main chromosome, but the majority of the genes associated with EPS production were on plasmid A. We additionally found candidate genes associated with nitrogen fixation across the main chromosome, including homologs of *nifH*, *nifDK*, *nifA*, *nifS*, *nifV*, *nifU* and *fixA* profiles (Figure S8), though other components of fixation were missing including *nifB*, *fixB*, and *fixX*.

### Genomic differentiation among treatments

Variant calling identified *de novo* variants in 91% of the derived strains (34 out of 37) when compared to the ancestor strain. These variants comprised: 1 large deletion (5869 bp in length), 11 instances of plasmid loss, and 129 single nucleotide polymorphisms (SNPs; see Data S2). The distribution of these variants was significantly structured by the selective nitrogen and salinity treatments (Table S6; Figure 4A). When examining the genetic distance between evolved strains, the selective nitrogen and salinity treatments significantly explained 5.5% and 5.7% of the variation, respectively. However, when excluding lost plasmids, there was only a marginal effect of the nitrogen treatment in predicting genetic distances. This is perhaps not surprising as plasmid loss was not evenly distributed among treatments, suggesting their loss underlies genetic differentiation between treatments (Figure 4B). Consistent with this, Fisher’s exact tests suggested Plasmid B was lost in the low-salinity treatment at a significantly higher rate than in the high-salinity treatment (Figure S9). Annotation of Plasmid B showed it to be enriched in four COG categories, including intracellular trafficking, cell motility, inorganic metabolism, as well as all DNA replication pathways. Plasmid C was lost only in the high salinity, low nitrogen treatments, in 40% of the strains of that treatment (Figure S9). Functions associated with defense, signal transduction, and an array of metabolic functions were enriched on this plasmid (Figure 4B).

**Figure 4:**
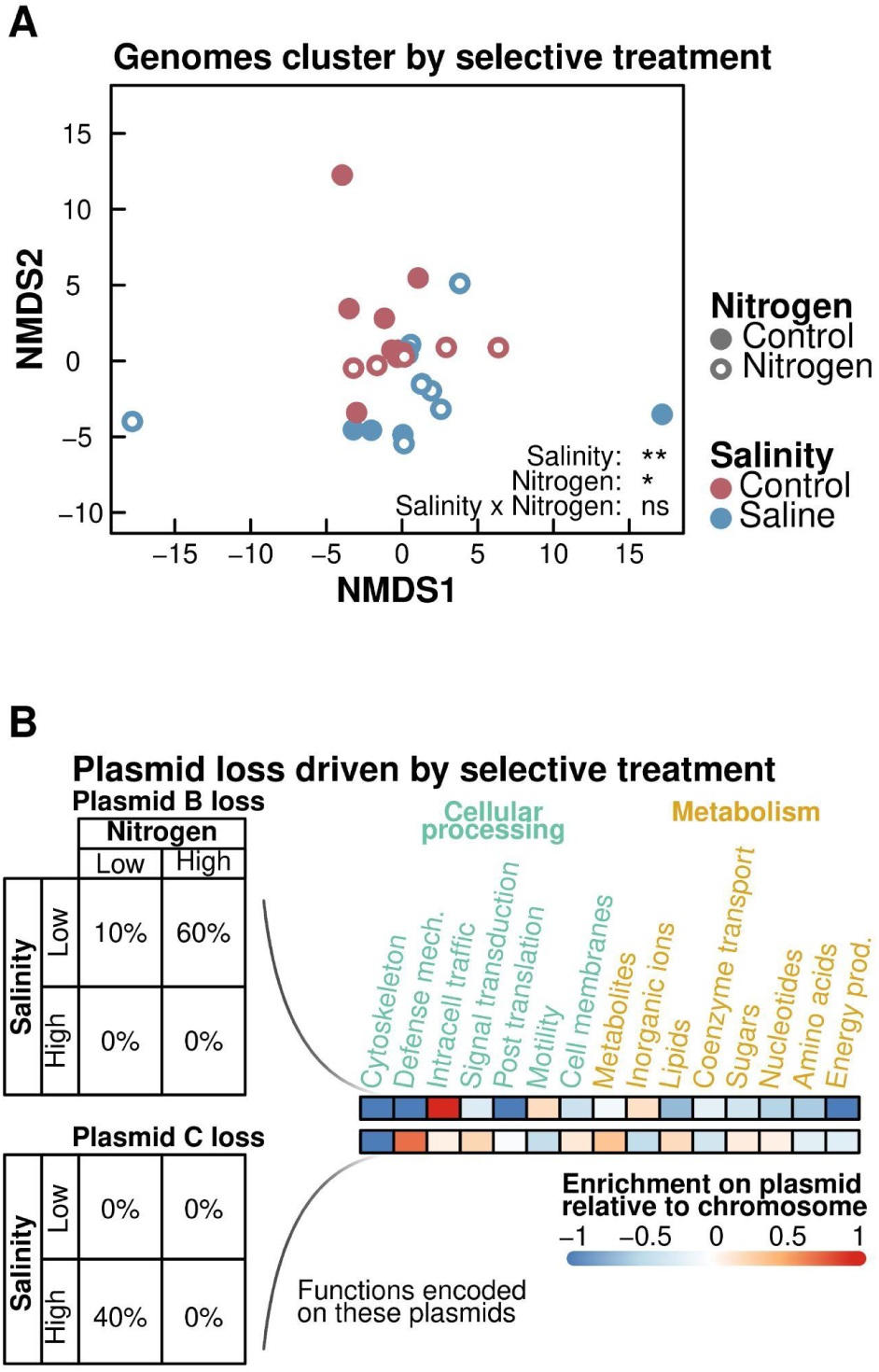
Selective treatments shape microbial genomes. Microbial genomes from the same selective treatment are more genetically similar (as measured by genetic distance) (**A**). Each point here represents a single microbial genome, and is coded by the strains’ evolutionary history: the historical salinity treatment (low vs. high salinity, delineated by blue vs. red coloring respectively) and the historical nitrogen treatment (low vs. high nitrogen, delineated by closed versus open circles respectively). The significance of these treatments is displayed in the bottom right of this panel. Full model results are in Table S5. We additionally display the **(B)** rate of loss of plasmids B and C across these selective treatments, as well as the enrichment of functions found on these plasmids compared to the main chromosome. Functions are broken down into major COG categories. For readability, we omitted functions associated with DNA replication, which are typically enriched on all plasmids.

SNPs were also not evenly distributed across treatments or replicons (Table S7), with the highest rate observed on the plasmids (Table S8). These SNPs were spread over 78 locations across the chromosome and plasmids, mapping to 46 unique genes. Of these, 45 had functional annotations. Multivariate models found that salinity, as well as its interaction with nitrogen, was a significant predictor of SNP distribution across functional annotation groups (Figure S10, Table S9). Individual assessment of these functional annotations found a significant increase in SNPs on genes annotated as phosphorelay signal transduction systems in the low-salinity treatments (Table S10). We additionally found a marginal increase in SNPs in high-nitrogen treatments on genes annotated as methytransferases, as well as a number of marginal interaction terms (see Table S11 for full details).

Many variants occurred on genes of high genomic differentiation, representing where mutations were consistently introduced on the exact same genes in the same selective treatment (Figures S11–S12, Table S12). Of particular interest may be an enrichment of SNPs on a nifS gene, a component of nitrogenase and driver of nitrogen fixation, in the high-nitrogen treatment, though this was seen most strongly in the high-salinity treatments. Additionally, the low-salinity treatments were enriched in SNPs across several genes associated with osmotic regulation, including membrane and biofilm regulation, as well as a solute-binding transport system.

## DISCUSSION

While beneficial plant-microbe interactions are often assumed to be the product of host selection on their microbial partners, our work demonstrates that host benefits can emerge strictly as a byproduct of microbial adaptation to abiotic stress in their free-living environment. Specifically, we found that at low nitrogen, *Allorhizobium* adapted to salt stress increased plant growth in high salinity, even though this microbial adaptation occurred entirely in the absence of a host. Moreover, across both salinity environments, we observed a strong positive correlation between microbial growth and plant benefits, where microbes with the highest partner quality in a given salinity environment were also the fastest-growing under those conditions. While this positive correlation between plant and microbial fitness, representing a fitness alignment between partners, might often be assumed to be the product of co-evolution, here it emerged independent of host selection. This correlation may suggest the traits underlying microbial salinity adaptation contribute to variation in microbial partner quality. These findings highlight that abiotic selective agents in the free-living environment can act as independent drivers in the evolution of host-microbe symbioses.

We hypothesize that microbial salinity adaptation increased microbial benefits to plants growing at high salinity by facilitating continued access to large populations of a beneficial microbe. In the high-salinity environment, high salt-adapted microbes achieved larger population sizes than low salt-adapted microbes and therefore may have been sufficiently numerous to continue to benefit the plant. The ancestral *Allorhizobium* strain was highly beneficial to plants at low nitrogen, and genomic characterization of this strain identified a number of potential benefits these microbes could provide to plants, including genes associated with increasing access to multiple resources, as well as the production of important plant hormones and vitamins. Maintaining access to microbial resources can be particularly beneficial to plant growth in stressful environments, such as under high salinity conditions, as resources such as nitrogen and phosphorus are essential in the synthesis of key plant hormones and antioxidants involved in the plant stress response [45–47].

The ancestral *Allorhizobium* isolate is closely related to other bacterial strains that fix nitrogen, suggesting it might increase plant growth at low nitrogen by fixing dinitrogen into a plant-available form. While nitrogen-fixing bacteria are best known within specialized, symbiotic partnerships, such as between rhizobia and legumes, free-living nitrogen fixation is thought to be the ancestral state of these symbioses [17]. As nitrogen fixation is an energetically expensive process [18], these free-living bacteria commonly colonize the carbon-rich zones surrounding a plant and can benefit plants through the leakage of excess fixed nitrogen [19, 20]. The ancestral *Allorhizobium* strain was isolated on nitrogen-free media, annotation identified genes involved in the nitrogen-fixing pathways, and its benefit to plants was largest under low nitrogen, consistent with N fixation capacity. We note, however, that growth on nitrogen-free media could reflect trace nitrogen in the media. Moreover, candidate N-fixing genes were scattered across the *Allorhizobium* genome, with several important genes involved in the nitrogen-fixation pathway being absent, while others could be paralogs. Without an acetylene reduction assay, we cannot be sure the *Allorhizobium* strain is a true diazotroph, and many other functions might contribute to its plant growth promotion.

A reduction in partner quality in high-nitrogen-adapted strains is consistent with prior work showing the evolution of less cooperative partners under nutrient enrichment [21, 22, 48]. Several non-exclusive mechanisms could explain this result. First, we tested the evolved strains on plants growing under low N conditions, so high nitrogen-adapted *Allorhizobium* strains may have been less beneficial because they were not well adapted to low N conditions; indeed, under low N, strains adapted to high N grew to lower densities than strains adapted to low N (Figure 2) and microbial growth rate was correlated with their quality as plant symbionts (Figure 3). Second, the functions that underlie the benefit of the ancestor may be costly to maintain and useful to the microbe only when nitrogen is scarce; high nitrogen could relax selection on these functions, reducing the microbe’s capacity to benefit the plant. For example, some high-nitrogen-adapted strains carried SNPs in a gene annotated as *nifS*, a component of the nitrogen fixation process. If this strain fixes nitrogen, these SNPs may be degrading the strain’s fixation capacity, and plant access to fixed nitrogen. There is prior work showing the degradation of N fixation in symbiotic *Rhizobium leguminasarum* strains under high nitrogen [21, 22, 48]. Third, these strains may actually be evolving traits involved in directly harming the host plant, as we note many of these nitrogen-adapted strains reduced plant growth below that of uninoculated controls. Regardless of mechanism however, our results show that adaptation to abiotic conditions in the free-living environment can drive the emergence of both beneficial and harmful microbial effects on hosts

The observed bacterial salinity adaptation may have resulted from plasmid loss or SNPs associated with bacterial osmotic regulation. Plasmids are frequently enriched in genes mediating adaptation to extreme environments, and are often quickly gained or lost as bacterial lineages adapt to changing environmental conditions, and also incorporate mutations more rapidly than the main chromosome [49, 50]. Consistent with this, many low-salinity adapted strains either lost plasmid B or were enriched in SNPs on plasmid B. This plasmid may be involved in osmotic regulation, being enriched in genes associated with intracellular trafficking, including bacterial secretion systems and ion pumps [51, 52]. Our ancestral strain was originally isolated from an aquatic habitat near downtown Toronto, which can be highly saline due to runoff from winter road salt [53]. Consequently, this plasmid may have aided the ancestor in these saline environments, and subsequently been lost when not needed in our experimental low saline environments. Additionally, strains adapted to low salinity were enriched in SNPs in bacterial signal transduction and biofilm regulation genes (Figure S8, S9). As transduction can increase osmotic regulation [54] while biofilms can protect bacterial cells from osmotic stress [55], accumulation of SNPs here may be degrading these strains’ capacity to tolerate salinity. We additionally note that a number of the high salinity-adapted strains lost plasmid C. Although the relation of this plasmid’s function to salinity tolerance is unclear, plasmids represent large DNA replication costs [50]. Perhaps this plasmid had utility in the natural freshwater and benign habitats we isolated these strains, while the high stress salinity treatments led to a genomic streamlining and loss of this plasmid.

While we have identified some potential genomic underpinnings of the observed bacterial phenotypes, there may be still other unidentified genetic bases for the phenotypic variation among our derived bacterial strains. With limited prior work or functional validation in this non-model, field-isolated *Allorhizobium* strain, many of the SNPs occurred in genes with unclear functions, making it difficult to determine the specific traits underlying the plant-microbe interaction. Additionally, our sequencing efforts identified only a small number of SNPs per strain, many of which were observed only once across replicates. These SNPs may underlie some of the observed phenotypes as traditional approaches rely on repeatable genomic changes across evolving lineages. However, epialleles could also be contributing here. Epigenome changes in bacteria occur relatively quickly, and can be vertically inherited [56], representing a rapid mechanism for microbial transgenerational stress tolerance that our sequencing would not capture. Consistent with this hypothesis, high nitrogen-adapted strains were marginally enriched in SNPs on genes annotated as methyltransferase, which may alter methylation rate. Future work may consider model systems with the capacity to investigate specific functional and transcriptional variation.

These results may directly impact interpretations of habitat-adapted symbiosis. While prior work has shown microbes from stressful habitats can confer habitat-specific stress tolerance on their hosts [17, 19, 57], these benefits are often interpreted as the result of host selection. Our results show that habitat-specific benefits can instead emerge from abiotic selection on microbes in the free-living environment alone, so the habitat specificity of a benefit is not, by itself, evidence that hosts shaped the microbial traits involved. We note, however, that this does not preclude the host from acting as a significant evolutionary driver in other contexts. Host microbiomes are diverse, and there are numerous examples of highly specialized, obligate interactions where microbes may be evolving with their host [58, 59]. Rather than viewing abiotic and biotic selection on these microbial symbionts as mutually exclusive, these processes may operate in tandem, with these byproduct traits being the jumping off point for more specialized symbioses [15, 60]. For example, abiotic selection in the free-living environment may generate reservoirs of pre-adapted, beneficial “byproduct” microbial traits, which hosts might subsequently select upon. Moreover, our approach here does not address how hosts acquire and retain microbes in natural, diverse communities. Even if microbial benefits to hosts evolve as byproducts of microbial adaptation to the abiotic environment, hosts may nonetheless evolve to preferentially recruit these beneficial strains, such that recruitment would not be incidental, even if the beneficial traits arose as byproducts.

Overall, our findings reveal that microbial traits beneficial to their hosts do not require host-mediated selection, but can evolve independently through abiotic selection in a free-living phase. Indeed we found both beneficial and harmful microbial interactions evolve in response to our selective treatments in free-living environments. Specifically, microbial salinity tolerance was associated with increases in microbial partner quality in saline environments. As these traits emerged independent of the plant, abiotic selective agents may significantly contribute to the evolution of the microbial traits that affect host growth. With many host-associated microbes also found free-living in the environment [1, 9, 61], these abiotic drivers may have significant consequences on the evolution of microbial partner quality. As we move forward, by identifying the specific traits that underlie host-microbe interactions and their selective drivers, including both the abiotic and biotic environments, we can better understand how host growth promotion evolves in microbes.

## ACKNOWLEDGEMENTS

We thank Frederickson lab members for thoughtful feedback on experimental design, analysis, and throughout the writing process. MEF received funding support from the Gordon and Betty Moore Foundation (Grants GBMF9536 [doi.org/10.37807/GBMF9356] and GBM10635 [doi.org/10.37807/GBMF10635]) and the Natural Sciences and Engineering Research Council of Canada (NSERC) Discovery Grant program (Grant RGPIN-2021-03711). KDR was supported through a postdoctoral fellowship from the Faculty of Arts and Science at the University of Toronto, and KB was supported by a University of Toronto Centre for Global Change Science Summer Internship.

## AUTHOR CONTRIBUTIONS

KDR and MEF conceived the original design of the experiment. KDR and KB carried out experimental work. KDR conducted the bioinformatic and statistical analyses with input from MEF. KDR wrote the first draft of the manuscript, and all authors contributed to review and editing.

## DATA AVAILABILITY

All sequence data and the ancestral genome assembly are available as an NCBI BioProject (accession no.: PRJNA1356529; https://www.ncbi.nlm.nih.gov/bioproject/PRJNA1356529). Phenotypic data and all associated code can be found at https://github.com/KevinDRicks/allorhizobium-salinity-adaptation. Strains used here are available upon request.

## SUPPLEMENTAL MATERIAL

**Figure S1:**
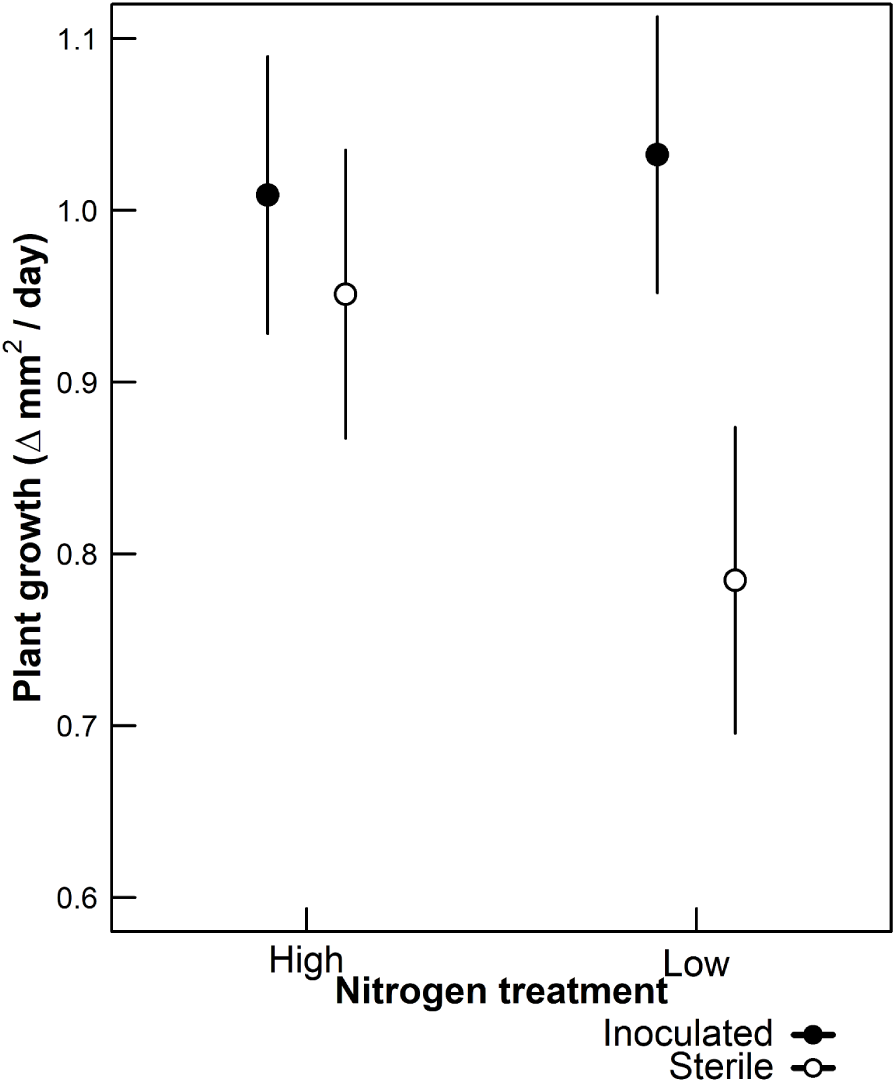
Plant growth (mm^2^/day) when inoculated with the ancestral *Allorhizobium* nitrogen-fixing strain (filled symbols) or uninoculated (open symbols) under low and high nitrogen. Plants in high nitrogen treatments were grown in Krazčič’s media, while the low nitrogen treatments were a Krazčič’s media modified to remove all nitrogen. Dots are estimated treatment marginal means. Bars are 95% confidence intervals generated from the standard error. Full model results are in Table S1.

**Figure S2:**
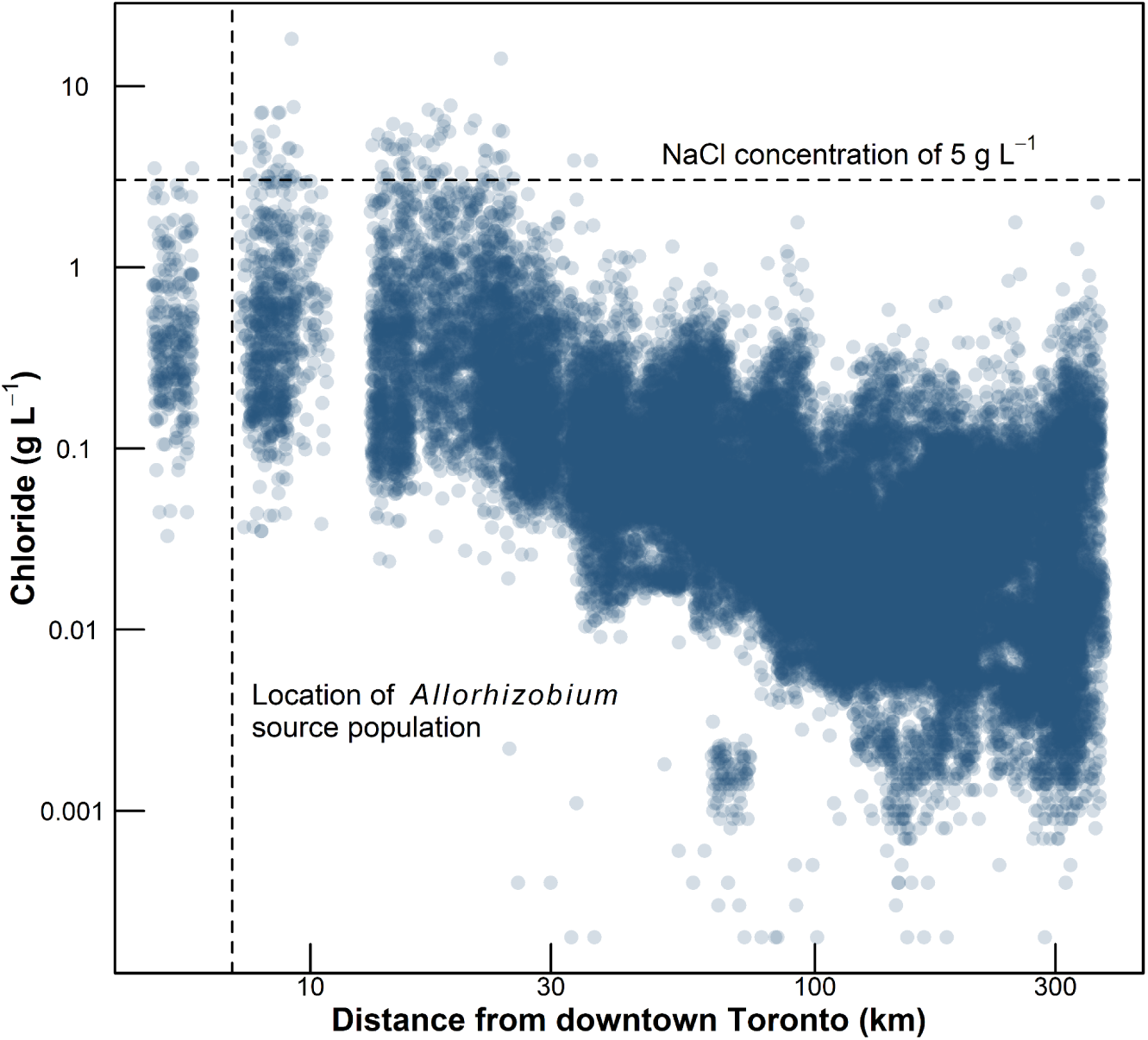
Chloride concentration of Ontario waterbodies increases closer to Toronto city center, the major metropolitan center in the region. Data are 48427 unique measurements from the Ontario Provincial Water Quality Monitoring Network as well as the Toronto and Region Conservation Authority, collected from 2010–2024 [1, 2]. The horizontal dotted line is the chloride concentration equivalent to 5 g L^-1^ NaCl (i.e., the high salinity treatment in our experiments). The vertical dotted line is the distance from downtown that our *Allorhizobium* strain was isolated from.

**Figure S3:**
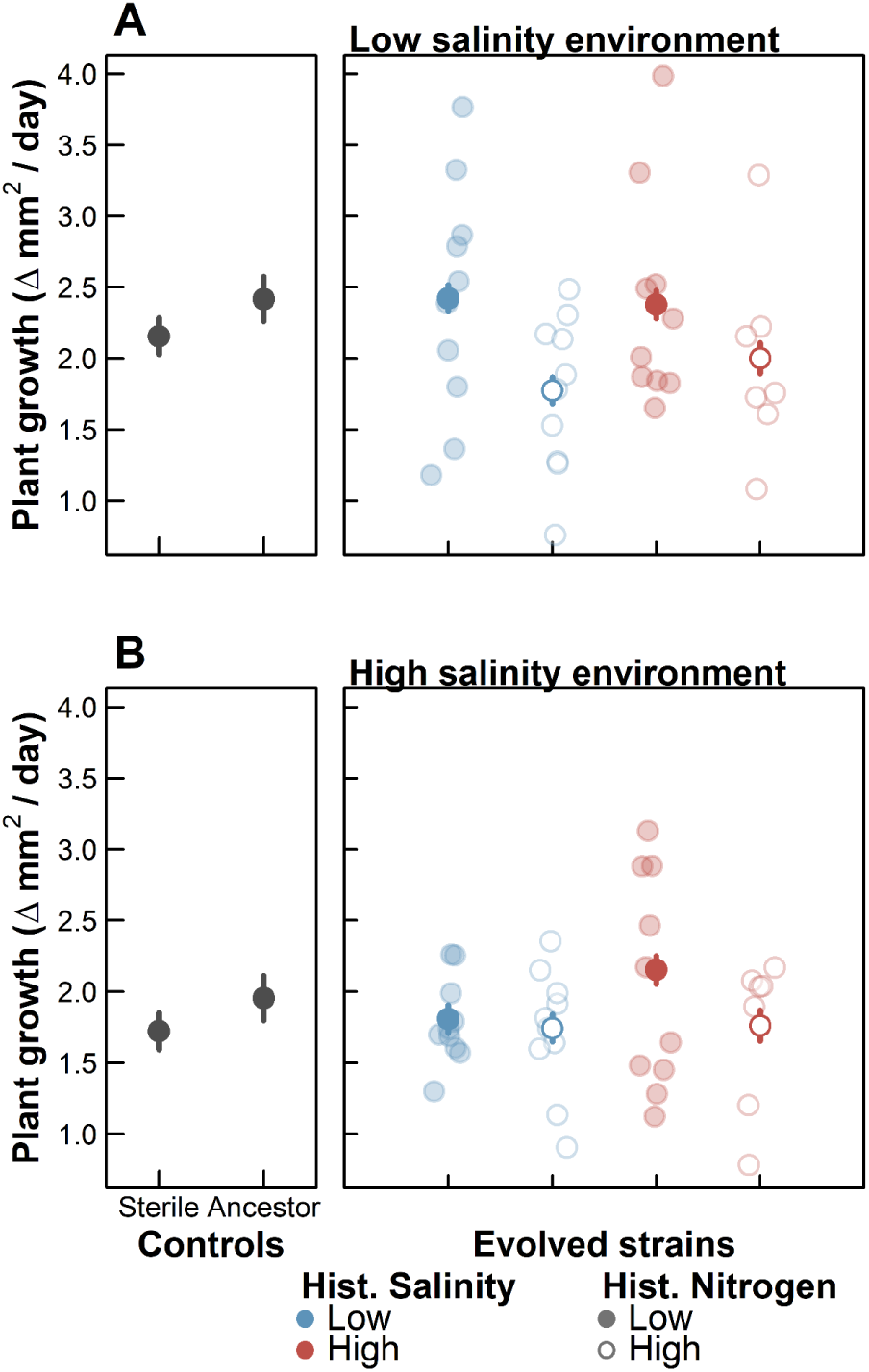
Evolutionary history of the experimentally evolved *Allorhizobium* strains impacts plant growth in both low (**A**) and high salinity (**B**) environments. Data are the same as in Figure 1, but show individual strain estimates. Dots are estimated marginal means of plant growth (mm^2^/day) when plants were inoculated with a sterile (i.e., microbe-free) control or the ancestral *Allorhizobium* (left panels, black symbols) or with *Allorhizobium* strains that evolved at low (blue symbols) or high (red symbols) salinity and low (closed symbols) or high (open symbols) nitrogen (right panels). Solid dots are estimated means by treatment (identical to Figure 1), while the semi-transparent dots represent the marginal means of individual evolved strains. Error bars are 95% confidence intervals generated from standard errors.

**Figure S4:**
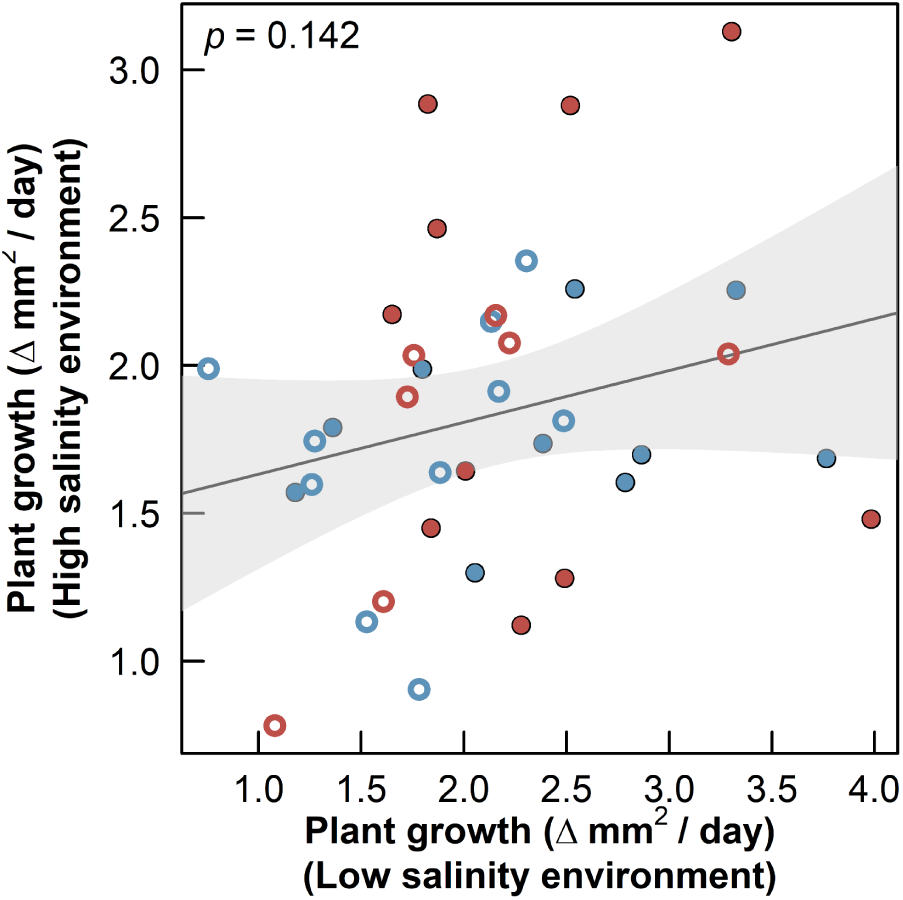
Microbial benefits provisioned to the host did not tradeoff between salinity environments. Dots are mean plant growth in high salinity (y-axis) and low salinity (x-axis) for individual *Allorhizobium* strains that evolved at low (blue symbols) or high (red symbols) salinity and low (closed symbols) or high (open symbols) nitrogen. Plant growth is the same as those presented in Figure 3. We include the predicted output from a linear regression correlating these traits, and the estimated *p* value in the top left corner of each panel.

**Figure S5:**
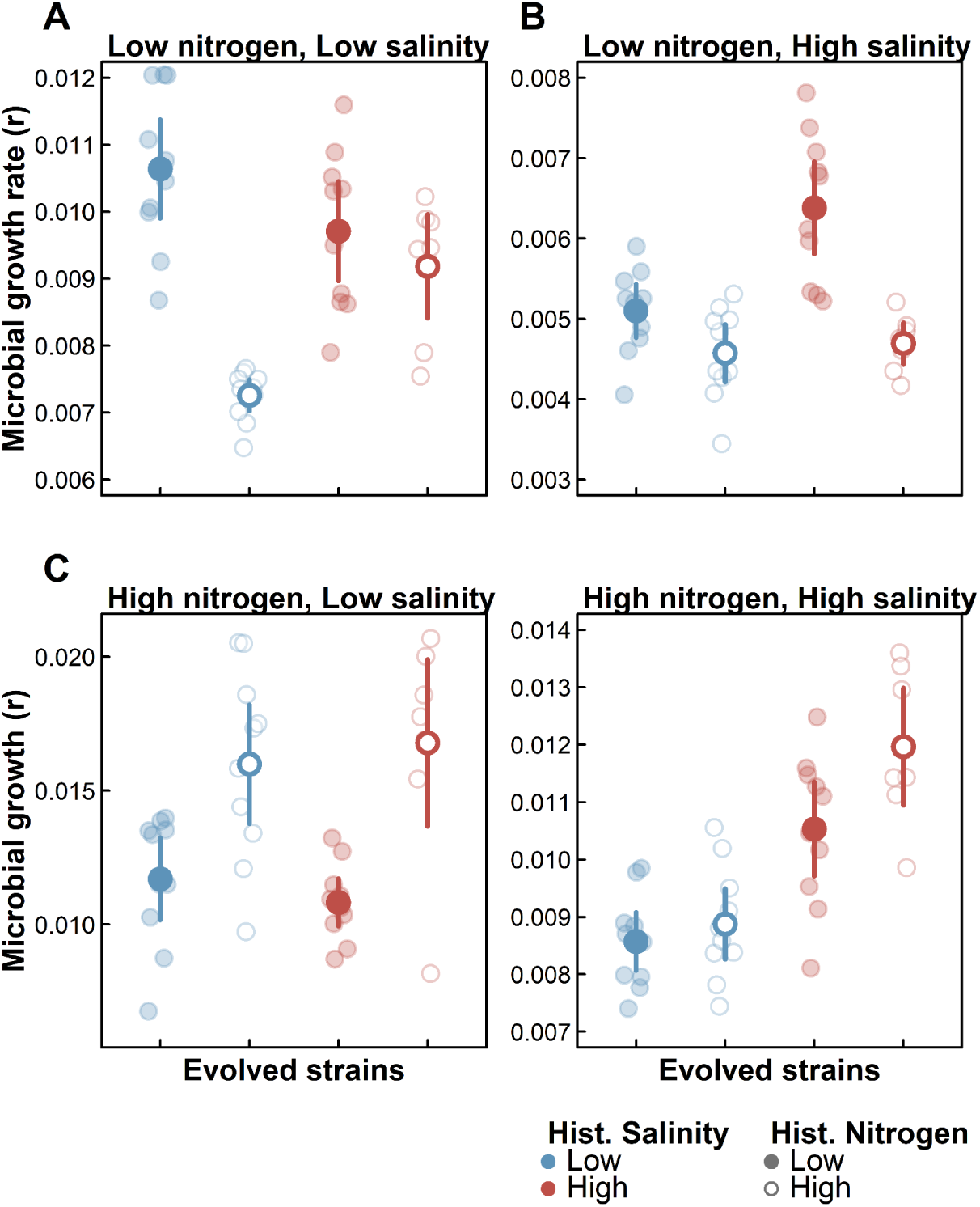
Evolutionary history of the experimentally evolved *Allorhizobium* strains impacts their growth rate across environments. Data are the same as in Figure 2, but show individual strain estimates. Solid dots are estimated marginal means of microbial growth (r, calculated from growth curves for each strain) for each selective treatment, while semi-transparent dots represent the marginal means for the individual strain estimates. Data are broken down by the contemporary environments: (**A**) low nitrogen and low salinity, (**B**) low nitrogen and high salinity, (**C**) high nitrogen and low salinity, and (**D**) high nitrogen and high salinity for *Allorhizobium* strains that evolved at low (blue symbols) or high (red symbols) salinity and low (closed symbols) or high (open symbols) nitrogen. Error bars are 95% confidence intervals generated from standard errors. Note the different y-axis scales between panels.

**Figure S6:**
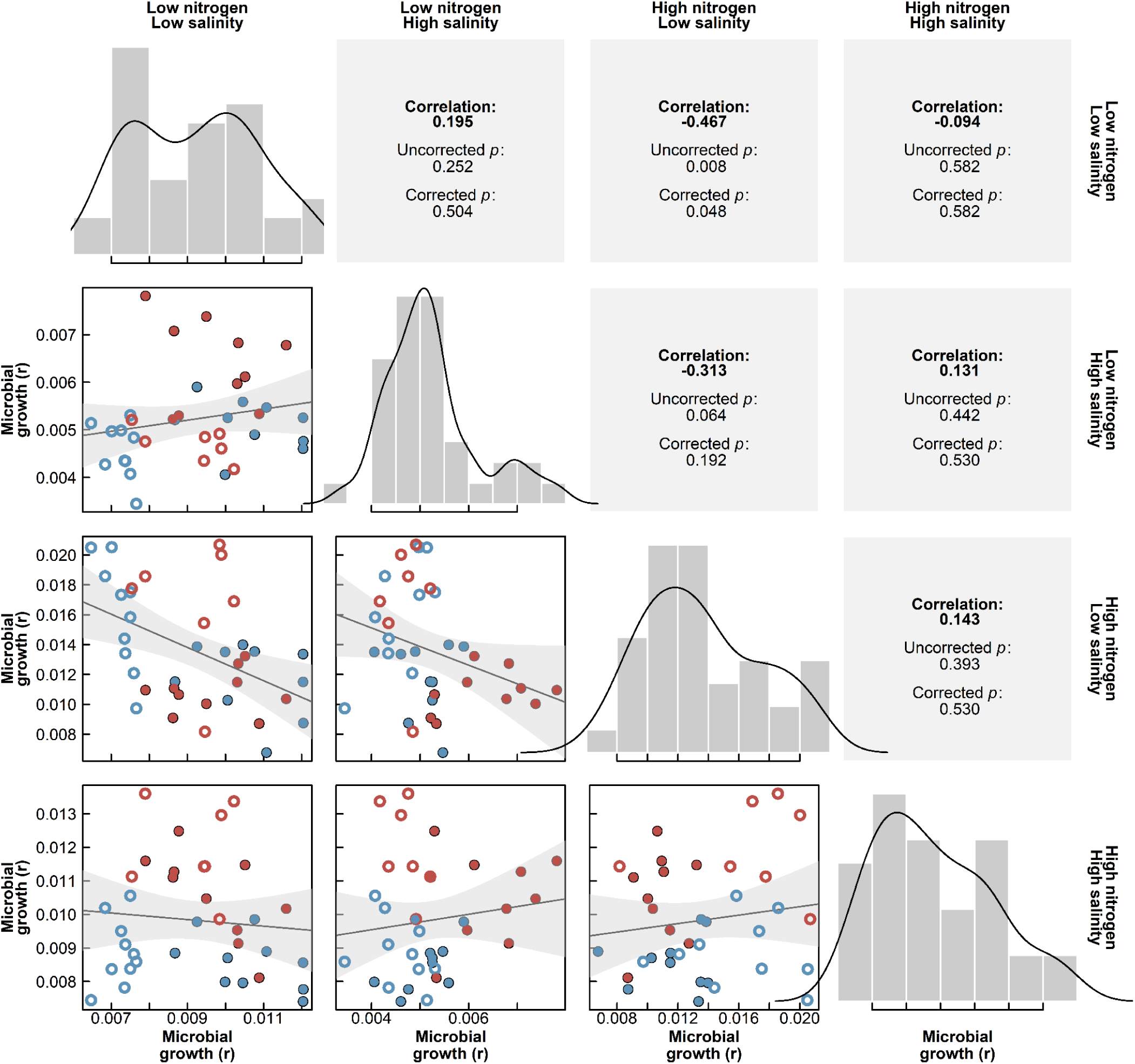
Tradeoffs in evolved microbial growth rate across all environments. The specific environment is coded in the top and right of the figure. The bottom left of the figure displays raw data, the top right displays the correlation coefficient for each pair, and the diagonal represents the distribution of each trait. Dots are each strain’s growth rate, coded by evolutionary history: low (blue symbols) or high (red symbols) salinity, and low (closed symbols) or high (open symbols) nitrogen. We display the line of best fit, as well as the 95% confidence interval in grey. We include the estimated *p* values for these correlations on the top right side, as well as *p* values corrected for multiple testing.

**Figure S7:**
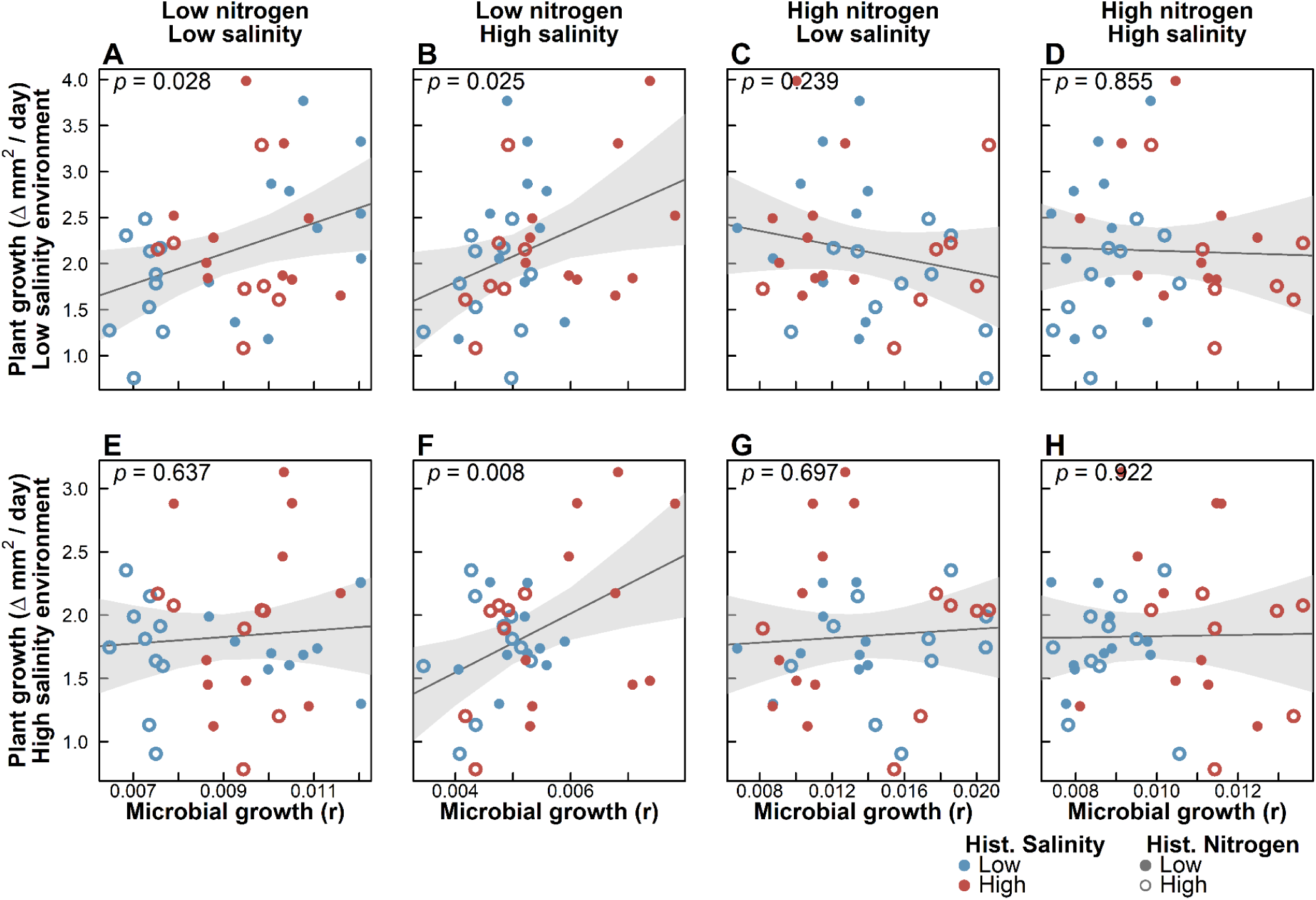
Microbial benefits provisioned to the host correlated with their growth across some of the salinity and nitrogen environments. Dots are mean plant growth rate (y-axis) and mean microbial growth rate (x-axis) for *Allorhizobium* strains that evolved at low (blue symbols) or high (red symbols) salinity and low (closed symbols) or high (open symbols) nitrogen. These comparisons are separated by plant growth in low salinity (**A-D**) and high salinity (**E-H**). This figure expands Figure 3 in the main text, which only showed panels **A** and **F**, which represent when the microbial growth environment matched the plant growth environment. We include the predicted output from a linear regression correlating these traits, and the estimated *p* value in the top left corner of each panel. Note the different y-axis scales in top and bottom panels.

**Figure S8:**
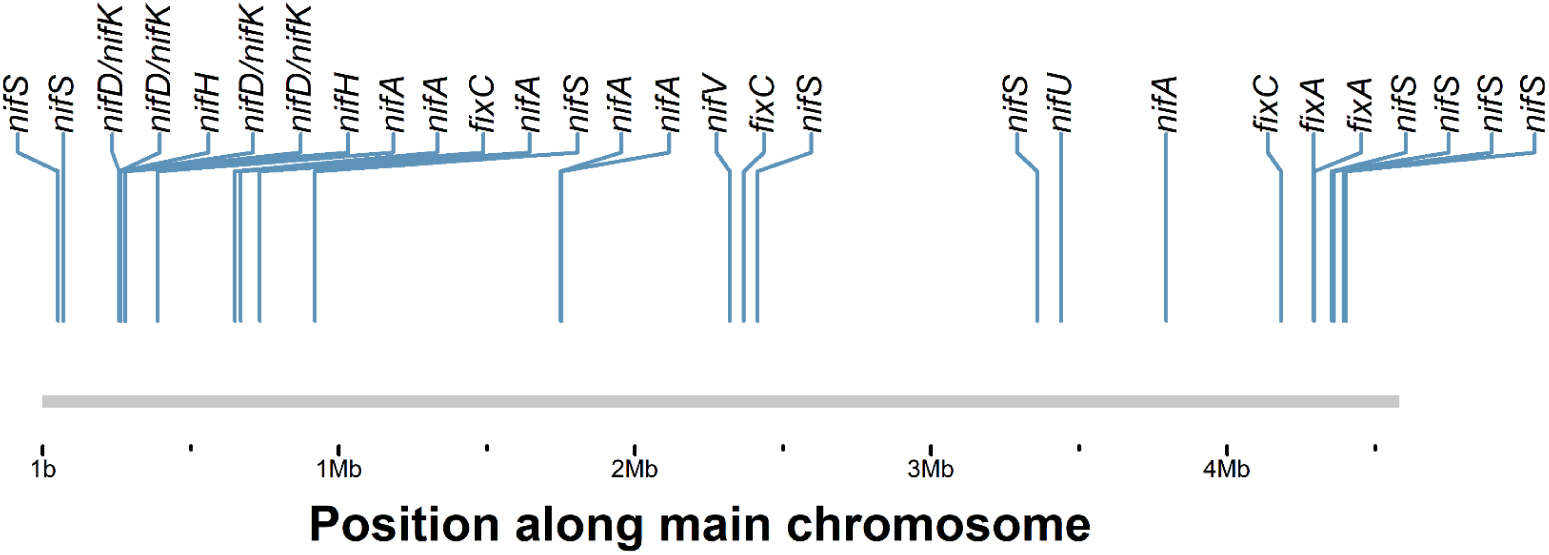
Distribution of candidate nitrogen fixation genes across the main chromosome. Genes were identified from Interpro protein family profiles using HMMER. Genes *nifD* and *nifK* have identical profiles and could therefore not be distinguished.

**Figure S9:**
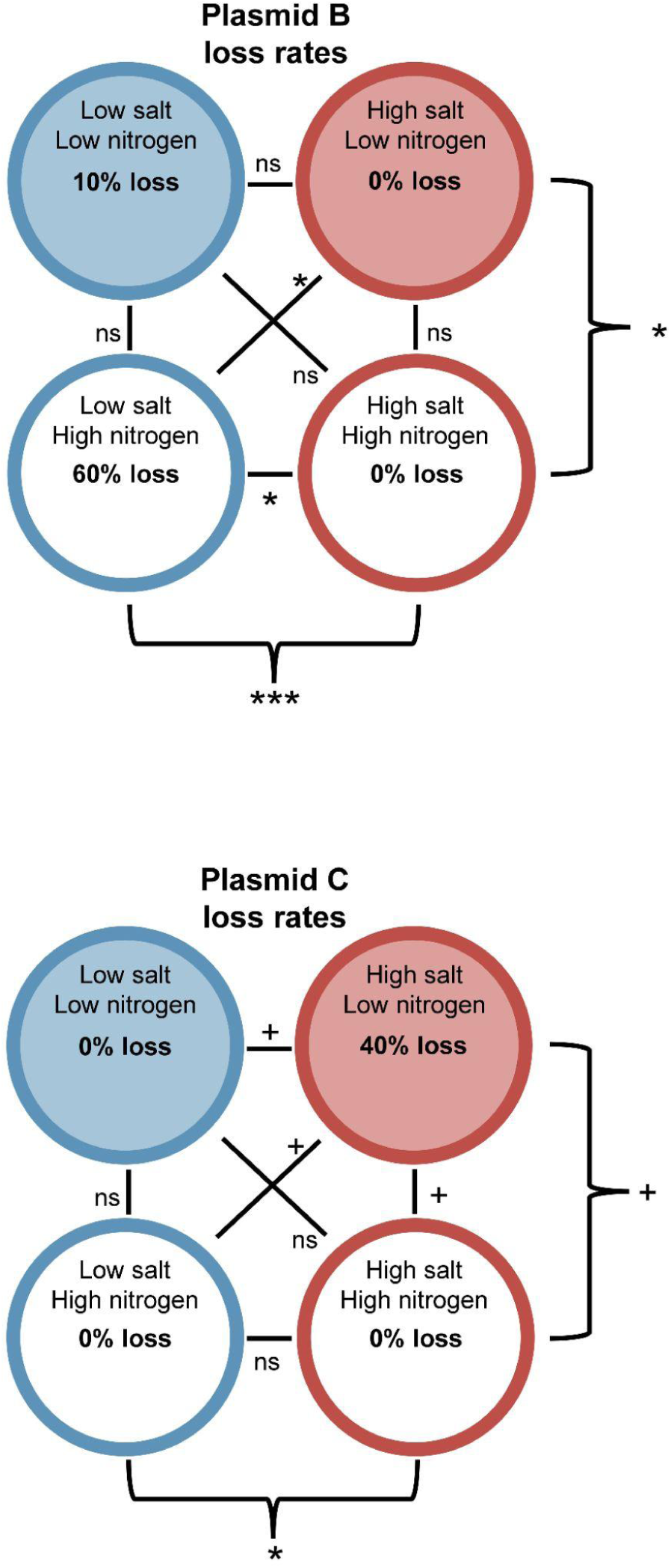
Rate of plasmid loss in evolved *Allorhizobium* strains by treatment. Differences in the rate of loss was compared between each treatment combination using Fisher’s Exact tests. The *p* values from each of these tests are summarized adjacent to each comparison as follows: ns p > 0.1; + p ≤ 0.1; * p ≤ 0.05; ** p ≤ 0.01; *** p ≤ 0.001

**Figure S10:**
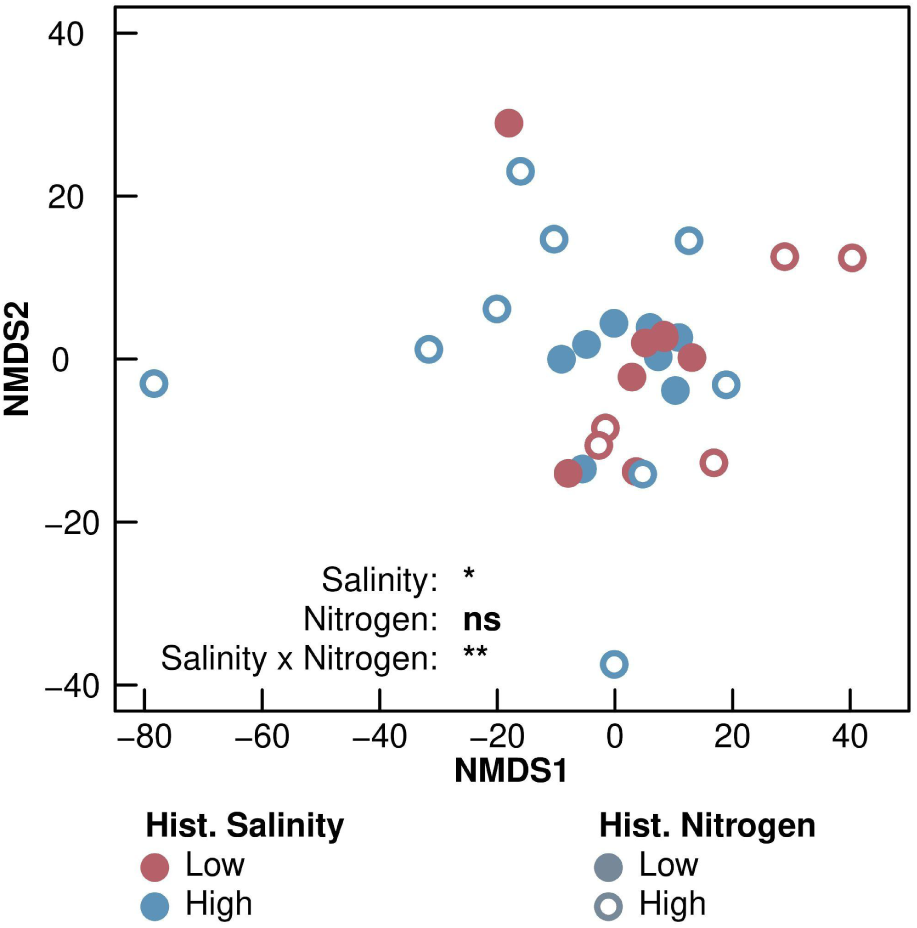
NMDS plot displaying distances between evolved genomes based on the annotations of their associated SNPs. Data are coded by the strains’ evolutionary history: the historical salinity treatment (low vs. high salinity, delineated by blue vs. red coloring respectively) and the historical nitrogen treatment (low vs. high nitrogen, delineated by closed versus open circles respectively. The significance of these treatments is displayed in the bottom right of this figure. Full model results are in Table S8.

**Figure S11:**
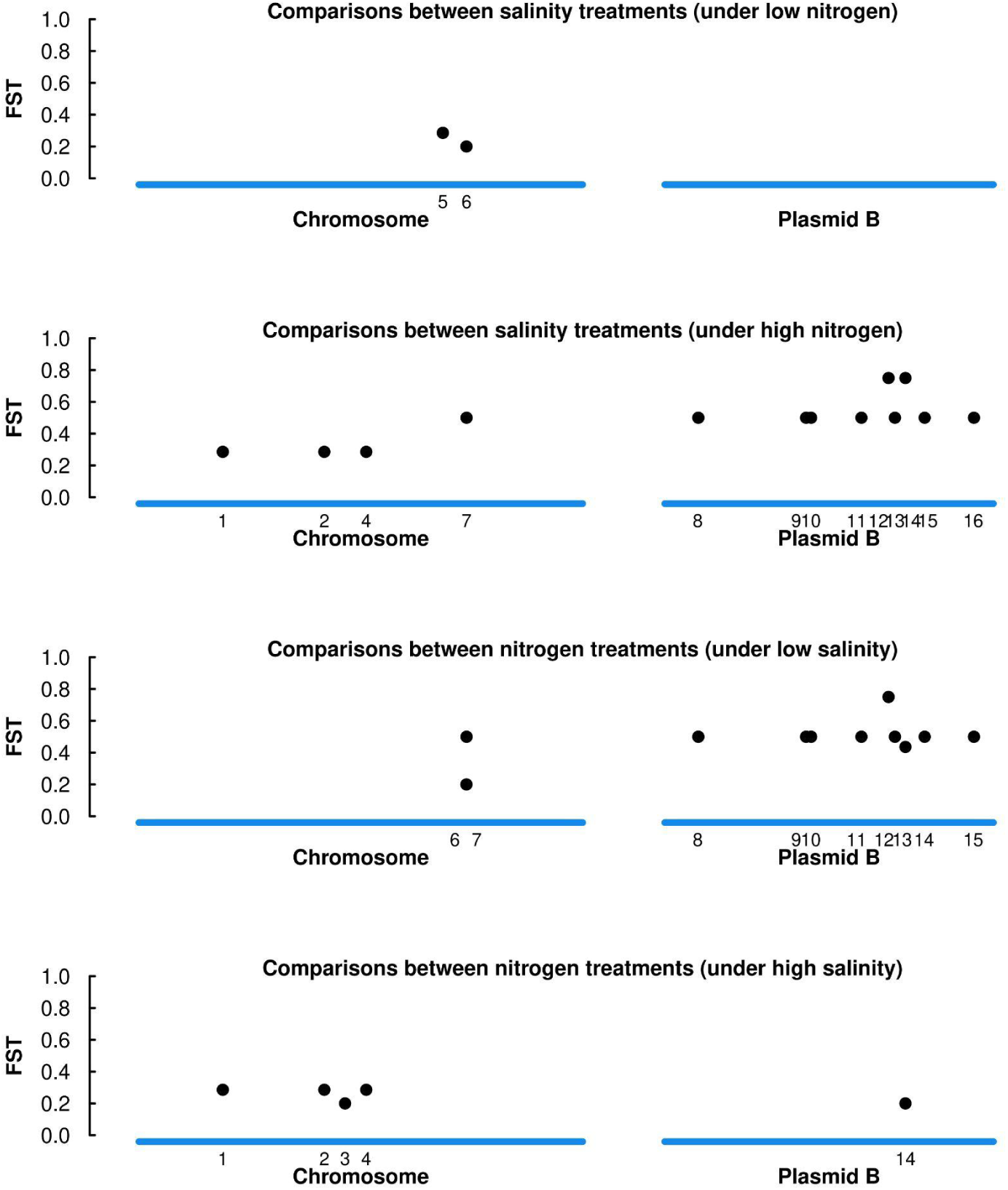
Pairwise differentiation in the enrichment of SNP on a gene by strain basis between the selective combinations. Differentiation here uses the same math as F_ST_, measuring the accumulation of SNPs on single genes. We specifically display only genes on the genome where differentiation in enrichment passed a 0.2 threshold. At the base of each panel, we include numbering to indicate which gene is represented at this site. Corresponding gene annotations are in Table S10. We display the enrichment of SNPs at each site for each treatment in Figure S9.

**Figure S12:**
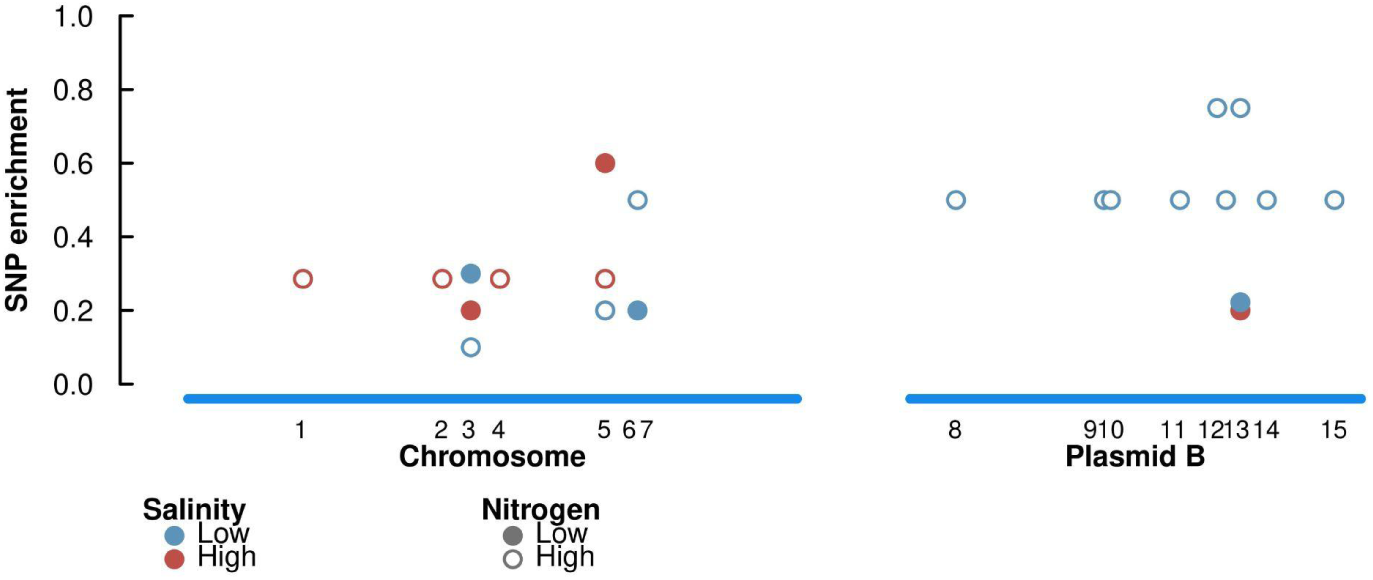
Enrichment of SNPs at each site across the genome, by selective treatment. These represent the same sites that pass the 0.2 threshold displayed in Figure S8. At the base of each panel, we include numbering to indicate which gene is represented at this site. Annotations of these genes can be seen in Table S10.

**Table S1:** Coefficient table characterizing plant growth with the ancestral *Allorhizobium* strain. Coefficients are extracted from a mixed effects model of plant area, measured over the course of the experiment. The model includes the following fixed effects: the timepoint when the image was taken, the microbial inoculum (inoculated with ancestral *Allorhizobium* vs. sterile), the contemporary nitrogen environment (low vs. high), and their interactions. We additionally included the starting size of plant material in the well, as well as its interaction with timepoint to control for differences in the starting size of plant material between wells. Individual well IDs and position in the growth chamber are random effects. Model coefficients here correspond with data displayed in Figure S1. Terms with a *p* value less than 0.1 are labelled in the Estimate column with the following: + *p* ≤ 0.1; * *p* ≤ 0.05; ** *p* ≤ 0.01; *** *p* ≤ 0.001.

| Term | Estimate | SE |  |
| --- | --- | --- | --- |
| Intercept | -0.928 | 1.939 |  |
| Inoculum (Sterile) | -0.890 | 2.043 |  |
| ContemporaryNitrogen (Low) | 2.517 | 2.038 |  |
| Starting Size | 2.769*** | 0.102 |  |
| Inoculum (Sterile) x ContemporaryNitrogen (Low) | -1.205 | 2.984 |  |
| Timepoint | 0.604*** | 0.037 | Terms representing plant growth rate |
| Timepoint x Inoculum (Sterile) | -0.086* | 0.039 |  |
| Timepoint x ContemporaryNitrogen (Low) | -0.005 | 0.037 |  |
| Timepoint x Starting Size | 0.042*** | 0.002 |  |
| Timepoint x Inoculum (Sterile) x Contemporary Nitrogen (Low) | -0.122* | 0.057 |  |

**Table S2:** Coefficient table characterizing plant growth with the evolved *Allorhizobium* strains. Coefficients are extracted from a mixed effects model of plant area, measured over the course of the experiment. The model includes the following fixed effects: the timepoint when the image was taken, the microbial inoculum’s historic nitrogen (low vs. high) and historic salinity environments (low vs. high), the contemporary salinity environment (low vs. high), and their interactions. We additionally included the starting size of plant material in the well, as well as its interaction with timepoint to control for differences in the starting size of plant material between wells. Strain ID, well ID, and position in the growth chamber are random effects. Model coefficients here correspond with data displayed in Figure 1. Terms with a *p* value less than 0.1 are labelled in the Estimate column with the following: + *p* ≤ 0.1; * *p* ≤ 0.05; ** *p* ≤ 0.01; *** *p* ≤ 0.001.

| Term | Estimate | SE |  |
| --- | --- | --- | --- |
| Intercept | -4.530* | 2.090 |  |
| ContemporarySalinity (Low) | 3.905* | 1.845 |  |
| HistoricSalinity (Low) | -1.099 | 1.737 |  |
| HistoricNitrogen (Low) | -2.856 | 1.825 |  |
| StartingSize | 3.148*** | 0.139 |  |
| ContemporarySalinity (Low) x HistoricSalinity (Low) | 0.885 | 2.407 |  |
| ContemporarySalinity (Low) x HistoricNitrogen (Low) | 0.539 | 2.460 |  |
| HistoricSalinity (Low) x HistoricNitrogen (Low) | 2.879 | 2.373 |  |
| ContemporarySalinity (Low) x HistoricSalinity (Low) x HistoricNitrogen (Low) | -2.805 | 3.289 |  |
| Timepoint | 1.155*** | 0.064 | Terms representing plant growth |
| Timepoint x ContemporarySalinity (Low) | 0.245*** | 0.067 |  |
| Timepoint x HistoricSalinity (Low) | -0.057 | 0.061 |  |
| Timepoint x StartingSize | 0.012* | 0.005 |  |
| Timepoint x HistoricNitrogen (Low) | 0.365*** | 0.063 |  |
| Timepoint x ContemporarySalinity (Low) x HistoricSalinity (Low) | -0.155+ | 0.087 |  |
| Timepoint x ContemporarySalinity (Low) x HistoricNitrogen (Low) | 0.001 | 0.089 |  |
| Timepoint x HistoricSalinity (Low) x HistoricNitrogen (Low) | -0.324*** | 0.084 |  |
| Timepoint x ContemporarySalinity (Low) x HistoricSalinity (Low) x HistoricNitrogen (Low) | 0.593*** | 0.119 |  |

**Table S3:** Coefficient table characterizing the microbial growth of the evolved *Allorhizobium* strains across variable nitrogen and salinity environments. Coefficients are extracted from a mixed model of microbial growth (r) estimated for each strain in 96-well plate growth assays. The model includes the following fixed effects: the microbial inoculum’s historic nitrogen (low vs. high) and historic salinity environments (low vs. high), the contemporary salinity environment (low vs. high), the contemporary nitrogen environment (low vs. high) and their interactions. Strain ID was a random effect. Model coefficients here correspond with data displayed in Figure 2. Terms with a *p* value less than 0.1 are labelled in the Estimate column with the following: + *p* ≤ 0.1; * *p* ≤ 0.05; ** *p* ≤ 0.01; *** *p* ≤ 0.001.

| Term | Estimate | SE |
| --- | --- | --- |
| Intercept | 0.012*** | 0.001 |
| ContemporaryNitrogen (Low) | -0.007*** | 0.001 |
| ContemporarySalinity (Low) | 0.005*** | 0.001 |
| HistoricNitrogen (Low) | -0.001 | 0.001 |
| HistoricSalinity (Low) | -0.003*** | 0.001 |
| ContemporaryNitrogen (Low) x ContemporarySalinity (Low) | 0.000 | 0.001 |
| ContemporaryNitrogen (Low) x HistoricNitrogen (Low) | 0.003** | 0.001 |
| ContemporarySalinity (Low) x HistoricNitrogen (Low) | -0.005*** | 0.001 |
| ContemporaryNitrogen (Low) x HistoricSalinity (Low) | 0.003* | 0.001 |
| ContemporarySalinity (Low) x HistoricSalinity (Low) | 0.002+ | 0.001 |
| HistoricNitrogen (Low) x HistoricSalinity (Low) | 0.001 | 0.001 |
| ContemporaryNitrogen (Low) x ContemporarySalinity (Low) x HistoricNitrogen (Low) | 0.003+ | 0.002 |
| ContemporaryNitrogen (Low) x ContemporarySalinity (Low) x HistoricSalinity (Low) | -0.004+ | 0.002 |
| ContemporaryNitrogen (Low) x HistoricNitrogen (Low) x HistoricSalinity (Low) | -0.002 | 0.002 |
| ContSalineLow x HistoricNitrogen (Low) x HistoricSalinity (Low) | 0.001 | 0.002 |
| ContemporaryNitrogen (Low) x ContemporarySalinity (Low) x HistoricNitrogen (Low) x HistoricSalinity (Low) | 0.003 | 0.002 |

**Table S4:**
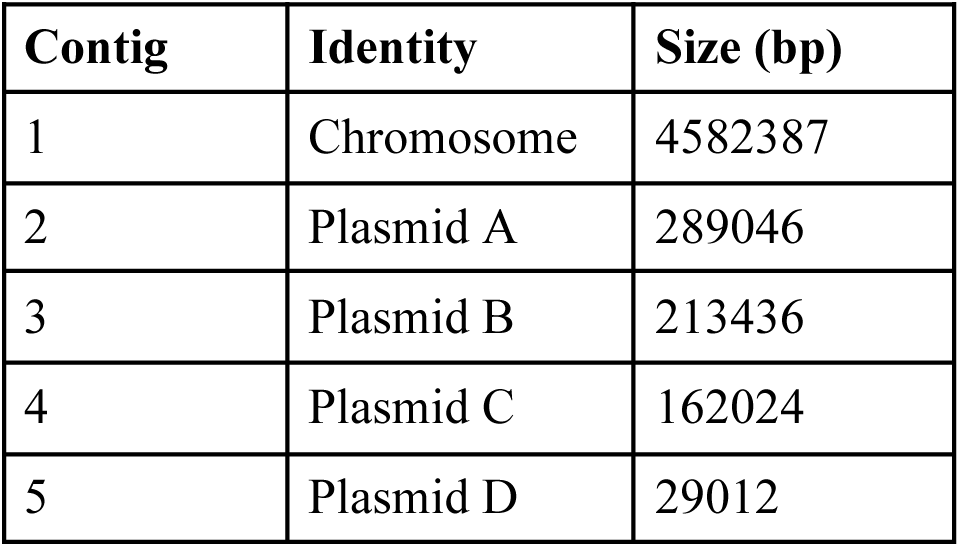
Summary of genome assembly of our ancestral strain of *Allorhizobium*, and the specific contigs therein.

**Table S5:** Distribution of microbial genes across our *Allorhizobium* genome with potential for plant-growth promotion. Genes were identified by comparing our *Allorhizobium* genome against the curated PLaBAse (PLant-associated BActeria) database. We breakdown genes across functions and genome location.

| Category | Function | Total | Main chromosome | Plasmid A | Plasmid B | Plasmid C | Plasmid D |
| --- | --- | --- | --- | --- | --- | --- | --- |
| Stress tolerance | EPS production | 15 | 6 | 9 | 0 | 0 | 0 |
| Nutrient acquisition | Siderophore production | 59 | 52 | 0 | 5 | 2 | 0 |
| Nutrient acquisition | Nitrogen fixation | 6 | 6 | 0 | 0 | 0 | 0 |
| Nutrient acquisition | Phosphate solubilization | 25 | 25 | 0 | 0 | 0 | 0 |
| Plant hormone production | Indole-3-acetic acid production | 17 | 17 | 0 | 0 | 0 | 0 |
| Plant hormone production | Cytokinin production | 7 | 7 | 0 | 0 | 0 | 0 |
| Plant hormone production | GABA production | 10 | 9 | 1 | 0 | 0 | 0 |
| Plant vitamin biosynthesis | Riboflavin biosynthesis | 13 | 12 | 0 | 0 | 1 | 0 |
| Plant vitamin biosynthesis | Thiamin biosynthesis | 7 | 6 | 1 | 0 | 0 | 0 |
| Plant vitamin biosynthesis | Pyridoxine biosynthesis | 9 | 9 | 0 | 0 | 0 | 0 |

**Table S6:**
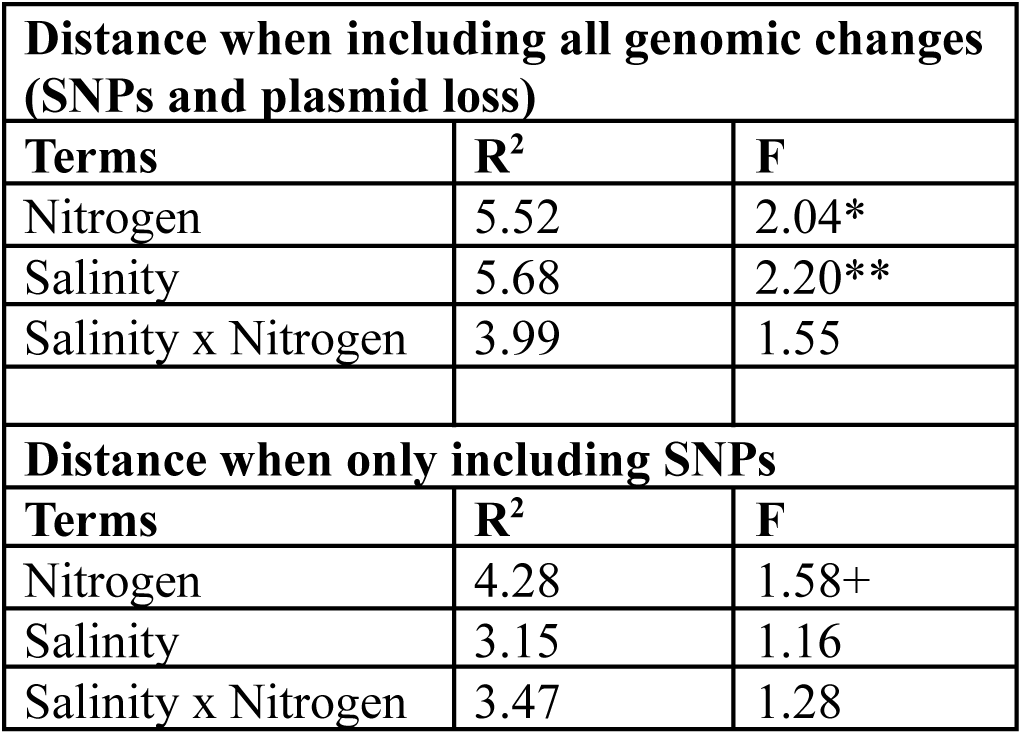
Result of PERMANOVA predicting genetic distance among evolved strains based on their selective treatment. We display results when plasmid loss is included (top) and when excluded (bottom). Figure 4A corresponds to the model including plasmids loss. For each term, we include the variance explained (R^2^) by that term as well as an *F* statistic. Terms with a p value less than 0.1 are indicated with the following: + *p* ≤ 0.1; * *p* ≤ 0.05; ** *p* ≤ 0.01; *** *p* ≤ 0.001.

| **Distance when including all genomic changes (SNPs and plasmid loss)** |  |  |
| --- | --- | --- |
| **Terms** | **$R^2$** | **F** |
| Nitrogen | 5.52 | 2.04* |
| Salinity | 5.68 | 2.20** |
| Salinity x Nitrogen | 3.99 | 1.55 |
| **Distance when only including SNPs** |  |  |
| **Terms** | **$R^2$** | **F** |
| Nitrogen | 4.28 | 1.58+ |
| Salinity | 3.15 | 1.16 |
| Salinity x Nitrogen | 3.47 | 1.28 |

**Table S7:** Average SNP count per strain, within a given treatment. The first line displays SNP count for the full bacterial genome, which are then broken down for each replicon.

|  | **Low Salinity** |  | **High Salinity** |  |
| --- | --- | --- | --- | --- |
|  | **Low Nitrogen** | **High Nitrogen** | **Low Nitrogen** | **High Nitrogen** |
| **Total** | 2.80 | 5.10 | 3.20 | 2.57 |
| **Chromosome** | 1.90 | 2.60 | 2.50 | 1.86 |
| **PlasmidA** | 0.0 | 0.10 | 0.20 | 0.57 |
| **PlasmidB** | 0.22 | 5.5 | 0.2 | 0.0 |
| **PlasmidC** | 0.6 | 0.2 | 0.5 | 0.14 |
| **PlasmidD** | 0.1 | 0.0 | 0.0 | 0.0 |

**Table S8:**
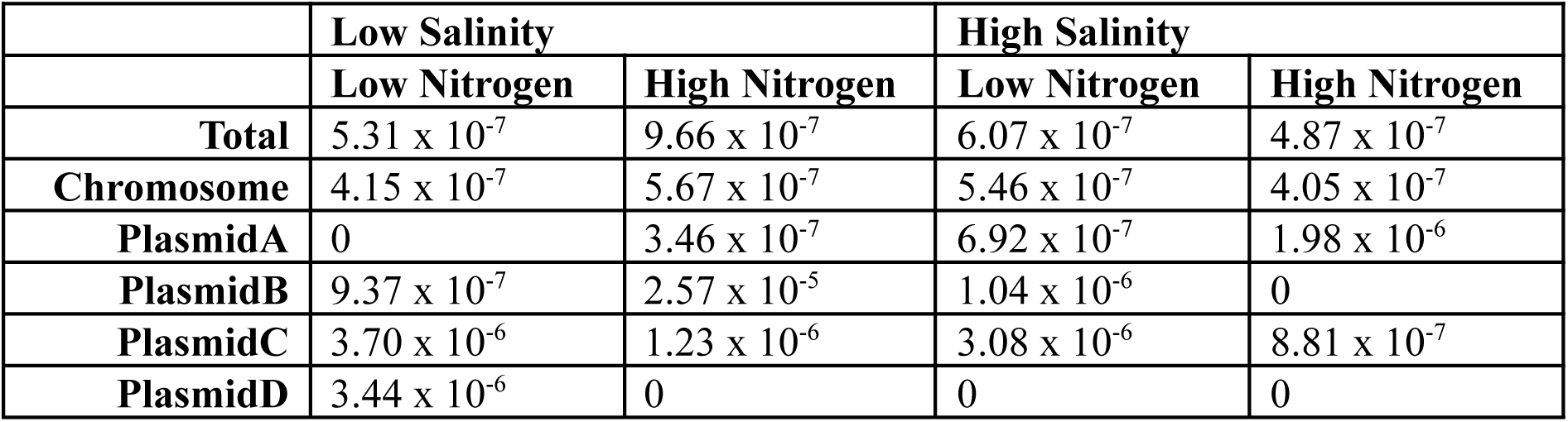
Average SNP frequency, when adjusting for the size of replicon, and replication, Data presented here are directly related to the frequencies observed in Table S6, with the SNP count divided by the size and number of replicons.

**Table S9:** Result of PERMANOVA predicting SNP distribution across functional annotation classes by their selective treatment. This analysis corresponds with the NMDS ordination displayed in Figure S7. For each term, we include the variance explained (R^2^) by that term as well as an *F* statistic. Terms with a p value less than 0.1 are indicated with the following: + *p* ≤ 0.1; * *p* ≤ 0.05; ** *p* ≤ 0.01; *** *p* ≤ 0.001.

| Terms | $R^2$ | F |
| --- | --- | --- |
| Salinity | 9.37% | 4.07* |
| Nitrogen | 2.19% | 0.95 |
| Salinity x Nitrogen | 12.74% | 5.55** |

**Table S10:** Classes of functional annotations with significantly different enrichment of SNPs between treatments. We display the specific functional class in the left column, and which treatments are enriched here.

| AnnotationClass | Low Salinity |  | High Salinity |  | Model output summary |
| --- | --- | --- | --- | --- | --- |
|  | Low Nitrogen | High Nitrogen | Low Nitrogen | High Nitrogen |  |
| phosphorelay signal transduction system | 0.2 | 0.5 | 0 | 0 | Significant effect of salinity |
| phosphorylation | 0 | 0.5 | 0.1 | 0 | Marginal interaction term |
| transferase activity, transferring phosphorus-containing groups | 0 | 0.5 | 0.1 | 0 | Marginal interaction term |
| phosphorelay sensor kinase activity | 0 | 0.5 | 0.1 | 0 | Marginal interaction term |
| signal transduction | 0 | 0.5 | 0.1 | 0 | Marginal interaction term |
| methyltransferase activity | 0 | 0.1 | 0 | 0.3 | Marginal effect of nitrogen |

**Table S11:** Genes with differential SNP enrichment values between treatment combinations. These correspond with the genes displayed in Figures S8 and S9. We additionally report the putative gene functions from both gene annotations as well as GO terms at each location.

| **F<sub>ST</sub> ID** | **Replicon** | **Annotation**<br>**(Eggnog)** | **Annotation**<br>**(Interpro GO Terms)** |
| --- | --- | --- | --- |
| 1 | Chromosome | Trimethylamine methyltransferase | Trimethylamine methyltransferase |
| 2 | Chromosome | nifS | No assigned function |
| 3 | Chromosome | No assigned function | No assigned function |
| 4 | Chromosome | Homocitrate synthase | Alpha-isopropylmalate synthase, Citramalate synthase, Pyruvate carboxyltransferase |
| 5 | Chromosome | No assigned function | Flagellin |
| 6 | Chromosome | Histidine Phosphotransfer domain | Signal transduction |
| 7 | Chromosome | Histidine Phosphotransfer domain | Signal transduction |
| 8 | PlasmidB | No assigned function | Lysozyme, Transglycosylase |
| 9 | PlasmidB | No assigned function | Phosphodiester glycosidase |
| 10 | PlasmidB | ABC-type sugar transport system | Solute-binding |
| 11 | PlasmidB | No assigned function | No assigned function |
| 12 | PlasmidB | No assigned function | No assigned function |
| 13 | PlasmidB | No assigned function | No assigned function |
| 14 | PlasmidB | No assigned function | Transposase |
| 15 | PlasmidB | Integrase | Integrase, Ribonuclease |
| 16 | PlasmidB | Diguanylate phosphodiesterase | Membrane regulation, Biofilm regulation |

## SUPPLEMENTAL METHODS

### Isolation and phenotyping of the ancestral *Allorhizobium* strain

In October 2023, we isolated bacterial strains from a population of duckweeds in a pond on the edge of downtown Toronto, Canada (Lat. 43.61990, Long. -79.33941). The site was a small pond filled with duckweed and other aquatic plants in Tommy Thompson Park, a headland that extends into Lake Ontario. We first gently rinsed collected duckweed in sterile saline solution (0.9% NaCl) to remove debris, and then vortexed them in fresh saline solution to detach bacterial cells from the plants. We spread these cells across solid Burk’s media (a nitrogen-free media commonly used for culturing nitrogen-fixing bacteria [3]), and picked and serially re-streaked single colonies on new plates. We taxonomically identified the isolates we obtained by amplifying and sequencing the 16S rRNA gene using the 27F and 1492R primers, and blasting sequences against the NCBI database[4]. We haphazardly chose one isolate from these strains with a full length *16S* sequence identified as belonging to the genus *Allorhizobium*.

In a pilot experiment, we evaluated the growth benefits of this *Allorhizobium* strain to *Lemna japonica*. We aseptically moved single fronds from a source population of duckweed (identical to that used in the main text, isolated from a ravine in Toronto: Lat. 43.68972, Long. -79.41944) into individual wells in sterile 24-well plates filled with Appenroth growth media. We filled each well with 2 mL of either Krazčič’s media [5], or a modified, nitrogen-free Krazčič’s media. In this nitrogen-free media, we replaced the nitrogen salt, KNO_3_, with equivalent concentrations of KCl to maintain potassium concentrations. We included 10 replicate wells inoculated with the *Allorhizobium* strain and 10 replicates inoculated with a sterile blank. We grew our *Allorhizobium* strain in liquid YM, pelleted it, and resuspended in nitrogen-free Krazčič’s media. We then added 75 μL of the strain to bring the optical density (OD_600_) of the solution to 0.015. We characterized plant growth over 18 days using identical methods as described in the main text and elsewhere [6] with our high-throughput imaging system. Using images captured over the course of this experiment to estimate proxies of plant fitness, we built parallel statistical models as to those described in the main text. From these models we extracted growth rate parameters for each treatment group (Figure S1, Table S1).

### Microbial sequencing and assembly

To prepare samples for whole genome sequencing, we streaked out freezer stocks of both our ancestral strain and all evolved strains on YMA. Single colonies were picked and grown to high density in liquid YM. Genomic DNA was then extracted using PureLink Genomic DNA extraction kits (Life Technologies) following the manufacturer’s recommended protocol.

We then used a combination of short and long-read sequencing to characterize these samples. Short-read sequencing for all strains was conducted at the Centre for the Analysis of Genome Evolution and Function (CAGEF, University of Toronto) using an Illumina NextSeq2000 XLEAP P1 sequencing system with paired 150 bp reads. All library prep was conducted by the CAGEF facility using Illumina DNA Prep Kits. Coverage was at minimum 75x, and ranged up to 300x. We additionally built a high-quality reference for the ancestral strain using long-read Oxford Nanopore Technologies (ONT) sequencing, completed by Plasmidsaurus (Eugene, OR). Briefly, amplification-free long read sequencing libraries were prepared using ONT v14 library prep chemistry. Sequencing was performed on ONT R10.4.1 flow cells. Raw data was processed using ONT’s basecalling model, outputting sequences with 150x genome coverage.

The ancestral long-read data was used to generate a *de novo* closed genome assembly using Trycycler [9]. Prior to assembly, raw long reads were filtered by length (>1 kbp) and quality (top 95%) using Filtlong [10]. The filtered reads were then subsampled into 12 independent subsets. Following the Trycycler pipeline, we used three different assembly methods to generate independent whole-genome assemblies, including Flye, Minimap2, and Raven [11–13]. Trycycler was then used to merge these assemblies into a final consensus genome. The consensus assembly was subsequently polished with Polypolish and Pypolca [14, 15] using the corresponding ancestral Illumina short-reads. This final polished assembly, along with the associated raw sequencing data, has been deposited as a component of an NCBI BioProject (https://www.ncbi.nlm.nih.gov/bioproject/PRJNA1356529).

### Evolutionary genomics

We broadly characterized if the distributions of variants were structured by the selection treatment. Using the ancestral genome as a scaffold, we overlaid variants onto the genome to create a full genome assembly for each evolved strain. We then aligned all genomes using the *DECIPHER* R package [7], and calculated Hamming distances between aligned genomes using the *pwalign* R package. We assessed if these distances were structured by treatment by piping the distance matrix into a permutational MANOVA (PERMANOVA) model, using the adonis2 function in the *vegan* R package [8], with their selective treatments as predictor variables. Plasmid loss, however, would overwhelm these analyses, representing an enormous deletion event. As plasmid loss represents an important component of the evolution of these strains, we recoded plasmid loss as a small deletion event at the end of the plasmid. We additionally conducted a parallel analysis removing all instance plasmid loss, to examine the impact of only SNPs on this analysis.

In evaluating plasmid loss across treatments, we first used Fisher’s exact test to conduct pairwise comparisons between treatments. Then, as plasmid loss represents a large deletion event, we characterized potential broad functional profiles of these plasmids using our annotations. The two plasmids lost in this experiment, plasmids B and C (see Results), represented significant functional diversity, with 221 and 120 unique gene annotations found on each, respectively. As characterizing the potential functions of these plasmids, given this diversity, is difficult to interpret, we binned genes into the COG categories provided by EggNog. We then characterized functional enrichment of each COG category on the plasmids by calculating a Sørensen–Dice coefficient for each category, subtracting the relative frequency of the function on the plasmid by the relative frequency of function on the chromosome, dividing by their sum [9]. Categories with >0 Sørensen–Dice coefficient may represent significant functions of the plasmid, relative to the main chromosome.

We examined how novel SNPs led to genomic differentiation by measuring pairwise SNP enrichment between treatments. We only considered SNPs here, as plasmid loss caused the entire plasmids to be highly differentiated. Given a scarcity of SNPs, we collapsed our data to a binary form, examining SNP occurence on a gene by strain basis: Any SNPs within a single gene would count as a single hit. We then calculated differentiation in SNP enrichment between treatments using F_ST_ calculations. This approach effectively asks whether rates of SNP accumulation on a given gene differ between treatments. We used the genet.dist function within the *hierfstat* R package [10] across the genome, with the haploid setting. Samples that were missing plasmids were removed from the analysis, as missing genes do not neatly code onto a mutation or the reference genome. Analogous to F_ST_, we focused specifically on sites with differentiation in SNP enrichment greater than 0.2.

We evaluated the distribution of SNPs based on their association with functional annotations using multivariate methods. We created a matrix representing the distribution of SNPs and their associated functions for each evolved strain: rows represented each strain and columns represented the observed functional term. This matrix was populated with 0s and 1s, with a 1 indicating the presence of a SNPs associated with that functional term for that strain. From this matrix we derived a distance matrix, based on Manhattan distances between each strain. We used this distance matrix in a PERMANOVA to evaluate similarity between treatments, with selective nitrogen and salinity treatments predictor variables. Additionally, we examined how individual annotation groups were associated with these treatments by building individual univariate models for each group. Models used binomial distributions, with historic nitrogen and salinity treatments as predictor variables, and Benjamini-Hochberg corrections for multiple testing [11]. Overall, our approach here is likely a conservative approach for evaluating the functional distribution of SNPs. Annotation pipelines are inherently imprecise, with a large fraction of the SNPs occurring in genes with no associated functions, which therefore had to be excluded from analysis.

